# A ligand-property-guided computational framework for prioritizing *de novo* protein binders for small molecules

**DOI:** 10.64898/2026.08.08.743643

**Authors:** Yaojun Zhu, Xiaoying Zhang

**Affiliations:** Chinese-German Joint Institute for Natural Product Research, Shaanxi International Cooperation Demonstration Base for Science and Technology, Shaanxi University of Technology, Hanzhong 723000, Shaanxi, China; Centre of Molecular & Environmental Biology, Department of Biology, University of Minho, Braga, Portugal

**Author notes:** **Corresponding author:** Dr. Xiaoying Zhang, *E-mail address:.

**Keywords:** Plant-derived small molecules, protein binder design, ligand-property- guided design, design consistency, *de novo* protein pocket design, capsaicin, (4R)- limonene, quercetin

## Abstract

Plant-derived small molecules possess highly diverse physicochemical properties, and the computational design of their protein recognition elements depends not only on the global structural quality of candidate backbones, but also on whether the local binding pocket, ligand-contact pattern, and predefined recognition conformation can be consistently retained after sequence design and structural back-prediction. To explore pocket-design strategies for different types of natural-product small molecules, this study selected capsaicin, (4R)-limonene, and quercetin as model ligands, representing a flexible amphipathic molecule, a compact hydrophobic monoterpene, and a rigid polyphenolic flavonoid scaffold, respectively, and covering the dimensions of pungent sensory flavor, volatile aroma, and flavonoid functional constituents. A ligand- physicochemical-property-guided computational design and multi-stage prioritization framework was established for candidate protein binders. The results showed that candidates with favorable initial global structural scores did not necessarily form reasonable local small-molecule binding pockets, indicating that evaluation of the local ligand environment is essential for candidate prioritization. After screening, 31 partial- pocket candidate backbones for capsaicin, 75 buried hydrophobic-pocket candidate backbones for (4R)-limonene, and 56 pocket-qualified candidate backbones for quercetin were obtained. Further sequence design and structural back-prediction analyses indicated that a subset of candidates could maintain the original pocket geometry and major ligand-contact patterns after sequence realization. Overall, these results suggest that the physicochemical properties of different plant-derived small molecules substantially influence the efficiency of de novo protein pocket formation, with compact hydrophobic ligands being more compatible with buried hydrophobic-pocket strategies, whereas flexible or multipolar ligands require a more refined balance between hydrophobic burial and polar exposure. This study provides a pre-experimental computational prioritization framework for natural-product small-molecule-recognizing proteins and offers candidate resources for subsequent protein expression, *in vitro* binding validation, active-constituent enrichment, and development of small-molecule biorecognition tools.

**Graphical Abstract:** 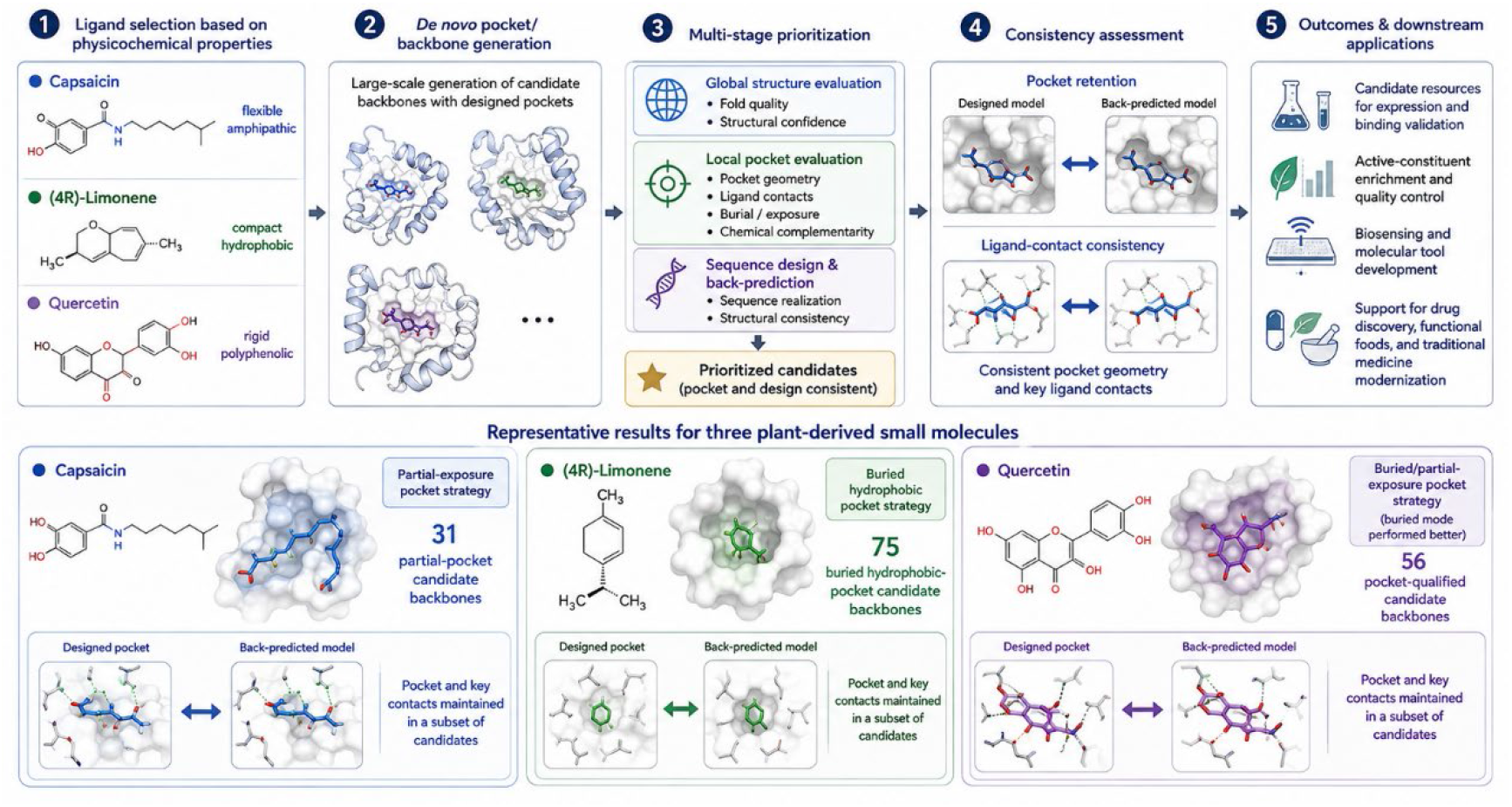

## 1 Introduction

Plant derived small molecules constitute an important chemical basis for drug discovery, functional food development, medicinal material quality control, flavor evaluation and the modernization of traditional medicine [1]. Recent advances in natural product research have increasingly emphasized the integration of chemical profiling, metabolomics, artificial intelligence and functional interpretation, which together expand the value of natural products beyond compound identification toward mechanism discovery and molecular tool development [2]. In addition to accurate qualitative and quantitative analysis, the development of programmable molecular recognition elements for plant derived small molecules is increasingly important for target capture, active constituent enrichment, biosensing and functional ingredient monitoring [3]. The Qinba region contains abundant medicinal and edible plant resources, and the quality, flavor and functional value of many regional products are closely associated with the composition and abundance of specific small molecule constituents [4]. Therefore, extensible recognition strategies for representative plant derived small molecules may provide useful molecular tools for the high value utilization and digital evaluation of characteristic natural resources.

However, protein-based recognition of natural-product small molecules remains challenging. Constrained by long-term evolution, natural proteins generally recognize endogenous molecules and common exogenous compounds associated with survival, signalling, and metabolism, whereas suitable natural receptors with sufficient affinity and specificity are often unavailable for non-natural, synthetic, or relatively rare plant-derived small molecules. Compared with proteins or peptides, small molecules have limited interaction surfaces, uneven polarity, variable conformational flexibility, and ligand-dependent requirements for burial or solvent exposure, making it difficult to form the broad, cooperative interfaces typical of protein–protein interactions. Recent studies of small-molecule-binding proteins and biosensors indicate that designed protein pockets can support ligand-responsive systems, but high affinity and specificity require precise pocket geometry, appropriately arranged polar and hydrophobic interactions, and rigorous structural and functional validation [5–7]. *De novo* design can address these limitations by controlling pocket geometry, electrostatic potential, and hydrophobic–hydrophilic balance at the atomic level and creating artificial microenvironments that are energetically and spatially complementary to specific ligands, thereby extending beyond the recognition and substrate spectra of natural proteins. It also enables large-scale computational screening before experimental validation, reducing reliance on animal immunization, associated ethical concerns and batch variation, and the dependence of conventional antibody generation on small-molecule immunogenicity and hapten conjugation, thus providing a systematic pre-experimental framework for developing stable, controllable, and rationally designed biorecognition tools for natural-product small molecules.

Capsaicin, (4R) limonene and quercetin provide a useful set of model ligands for evaluating such a design strategy because they differ substantially in hydrophobicity, polarity, rigidity, flexibility and functional context. Capsaicin is a flexible amphipathic molecule containing a vanillyl aromatic group, polar functional groups and a hydrophobic aliphatic chain, and is closely associated with pungent sensory quality in chili products [8]. (4R) limonene is a compact and highly hydrophobic monoterpene hydrocarbon representing volatile aroma related constituents in citrus resources [9]. Quercetin is a rigid polyphenolic flavonol scaffold with multiple hydroxyl groups and a carbonyl group, and is widely used as a representative flavonoid in phytochemical analysis and functional constituent studies [10]. These three ligands therefore cover distinct physicochemical and application dimensions, including pungent flavor, volatile aroma and flavonoid bioactivity, and allow comparison of different pocket conditioning strategies for chemically diverse plant derived small molecules.

In this study, capsaicin, (4R) limonene and quercetin were selected as model plant derived small molecules to establish a ligand property guided and design consistency guided computational framework for candidate protein binder prioritization. RFdiffusion based backbone generation, local pocket evaluation, LigandMPNN sequence design [14], RoseTTAFold3 structural back prediction [15], projected contact recovery analysis [16], molecular docking [17] and representative molecular dynamics simulation [18] were integrated to identify candidates that not only form plausible local binding pockets, but also retain the intended ligand recognition geometry after sequence realization. The resulting candidates should be regarded as pre experimental resources for subsequent protein expression, *in vitro* binding validation and development of natural product small molecule recognition tools.

## 2 Methods and materials

### 2.1 Study design and screening workflow

This study used a staged computational screening workflow to prioritize candidate protein binders for capsaicin, (4R)-limonene and quercetin. The workflow included ligand preparation, ligand-property-guided exposure conditioning, RFdiffusion3-based backbone generation, JSON-derived structural filtering, ligand-centred pocket evaluation, LigandMPNN-based sequence design, RoseTTAFold3-based structural back-prediction, projected ligand-pose compatibility analysis, sequence-level property assessment, integrated representative selection, molecular docking and representative molecular dynamics analysis.

At each stage, candidates were either retained in the principal analysis cohort, assigned to a conditional or repair-oriented cohort, or excluded according to predefined structural, geometric, sequence-level or ligand-compatibility criteria. The staged inclusion and exclusion logic is summarized in Table 1.

**Table 1.**
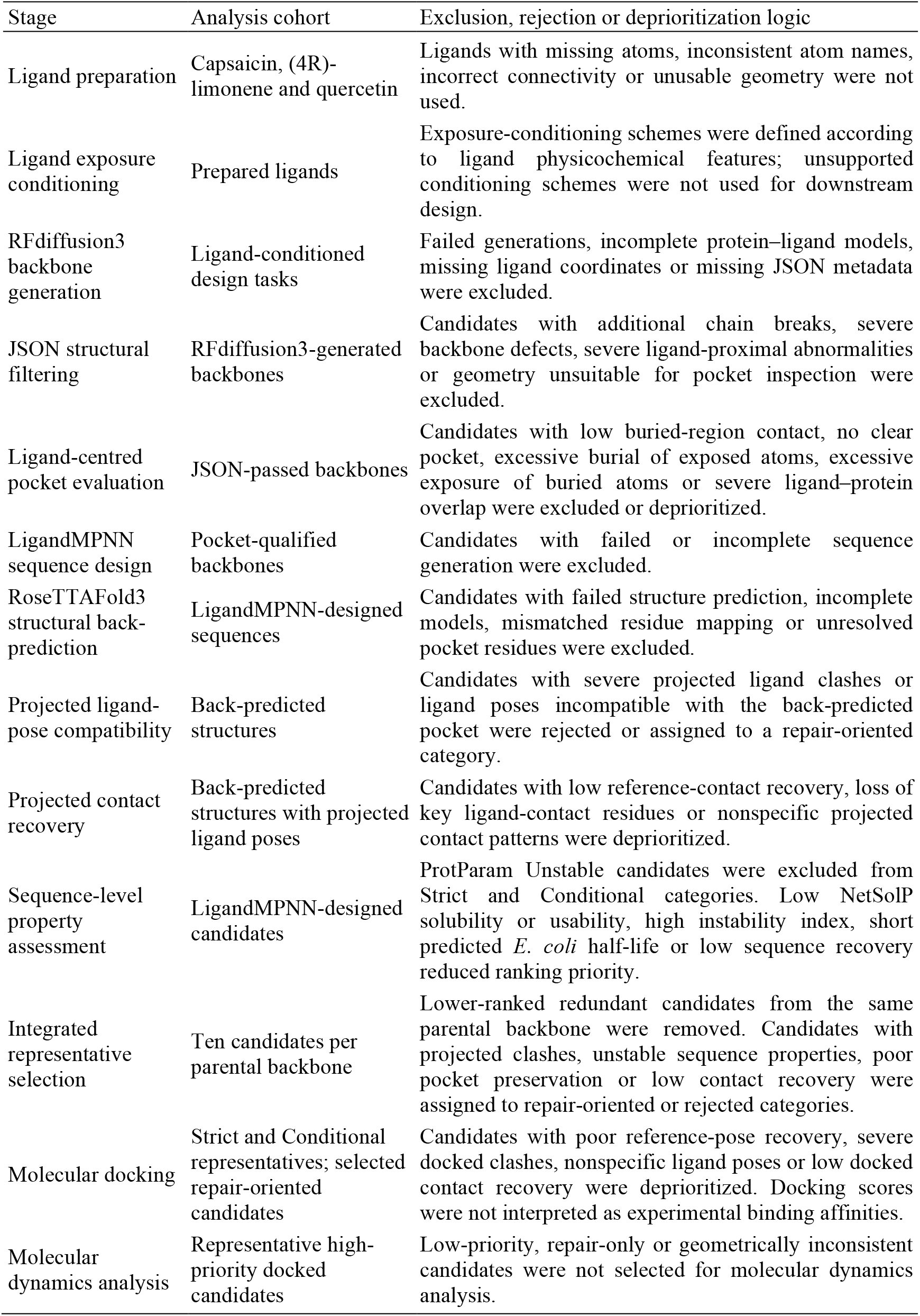
Staged inclusion and exclusion criteria used in the computational screening workflow.

| Stage | Analysis cohort | Exclusion, rejection or deprioritization logic |
| --- | --- | --- |
| Ligand preparation | Capsaicin, (4R)-limonene and quercetin | Ligands with missing atoms, inconsistent atom names, incorrect connectivity or unusable geometry were not used. |
| Ligand exposure conditioning | Prepared ligands | Exposure-conditioning schemes were defined according to ligand physicochemical features; unsupported conditioning schemes were not used for downstream design. |
| RFdiffusion3 backbone generation | Ligand-conditioned design tasks | Failed generations, incomplete protein–ligand models, missing ligand coordinates or missing JSON metadata were excluded. |
| JSON structural filtering | RFdiffusion3-generated backbones | Candidates with additional chain breaks, severe backbone defects, severe ligand-proximal abnormalities or geometry unsuitable for pocket inspection were excluded. |
| Ligand-centred pocket evaluation | JSON-passed backbones | Candidates with low buried-region contact, no clear pocket, excessive burial of exposed atoms, excessive exposure of buried atoms or severe ligand–protein overlap were excluded or deprioritized. |
| LigandMPNN sequence design | Pocket-qualified backbones | Candidates with failed or incomplete sequence generation were excluded. |
| RoseTTAFold3 structural back-prediction | LigandMPNN-designed sequences | Candidates with failed structure prediction, incomplete models, mismatched residue mapping or unresolved pocket residues were excluded. |
| Projected ligand-pose compatibility | Back-predicted structures | Candidates with severe projected ligand clashes or ligand poses incompatible with the back-predicted pocket were rejected or assigned to a repair-oriented category. |
| Projected contact recovery | Back-predicted structures with projected ligand poses | Candidates with low reference-contact recovery, loss of key ligand-contact residues or nonspecific projected contact patterns were deprioritized. |
| Sequence-level property assessment | LigandMPNN-designed candidates | ProtParam Unstable candidates were excluded from Strict and Conditional categories. Low NetSolP solubility or usability, high instability index, short predicted *E. coli* half-life or low sequence recovery reduced ranking priority. |
| Integrated representative selection | Ten candidates per parental backbone | Lower-ranked redundant candidates from the same parental backbone were removed. Candidates with projected clashes, unstable sequence properties, poor pocket preservation or low contact recovery were assigned to repair-oriented or rejected categories. |
| Molecular docking | Strict and Conditional representatives; selected repair-oriented candidates | Candidates with poor reference-pose recovery, severe docked clashes, nonspecific ligand poses or low docked contact recovery were deprioritized. Docking scores were not interpreted as experimental binding affinities. |
| Molecular dynamics analysis | Representative high-priority docked candidates | Low-priority, repair-only or geometrically inconsistent candidates were not selected for molecular dynamics analysis. |

### 2.2 Ligand structure retrieval and preparation

Three-dimensional structures of capsaicin, (4R)-limonene and quercetin were obtained from the wwPDB Chemical Component Dictionary (CCD) [19]. The CCD identifiers were 4DY for capsaicin, 9IR for (4R)-limonene and QUE for quercetin. Ligand structures were downloaded in CIF and SDF formats and inspected for atom identity, bond connectivity, stereochemical consistency, formal charge assignment and three-dimensional geometry. PyMOL 3.0.3 was used to confirm residue names, atom names and spatial integrity. The same ligand identities, atom names, protonation states and tautomeric states were retained throughout design, docking and downstream structural analyses.

Ligand exposure-conditioning schemes were defined according to ligand physicochemical features and intended pocket geometry. Capsaicin was treated as a flexible amphipathic ligand. Its aromatic scaffold and hydrophobic chain were assigned as buried or partially buried recognition regions, whereas polar oxygen- and nitrogen-containing groups were retained as partially exposed regions. (4R)-limonene was treated as a compact hydrophobic ligand, and its nonpolar carbon skeleton was assigned as the buried recognition region. Quercetin was evaluated under two conditioning schemes. In the partial-exposure scheme, the aromatic flavonol core was assigned as the main buried region and selected hydroxyl-containing regions were retained as exposed or partially exposed. In the buried scheme, the quercetin scaffold was assigned for more extensive enclosure within the designed pocket (Table 2).

**Table 2.** Ligand features and exposure-conditioning schemes.

| Ligand | CCD ID | Physicochemical category | Conditioning scheme | Buried or partially buried region | Exposed or partially exposed region | Purpose |
| --- | --- | --- | --- | --- | --- | --- |
| Capsaicin | 4DY | Flexible amphipathic ligand | Partial exposure | Aromatic scaffold and hydrophobic chain | Phenolic, methoxy and amide-associated polar atoms | Balance hydrophobic enclosure and polar accessibility |
| (4R)-limonene | 9IR | Compact hydrophobic monoterpene | Buried ligand | Nonpolar carbon skeleton | Not separately defined | Generate compact hydrophobic cavities |
| Quercetin | QUE | Rigid polyphenolic flavonol | Partial exposure | Aromatic flavonol core | Selected hydroxyl-containing regions | Test partial exposure of polar groups |
| Quercetin | QUE | Rigid polyphenolic flavonol | Buried ligand | Larger fraction of the flavonol scaffold | Not separately defined | Test extensive scaffold enclosure |

### 2.3 RFdiffusion3-based backbone generation and initial filtering

Candidate ligand-binding backbones were generated using RFdiffusion3 [20] with ligand structures supplied as non-protein conditioning components. Four independent design tasks were performed. Capsaicin was processed under the partial-exposure scheme with a binder length range of 100–150 residues. (4R)-limonene was processed under the buried-ligand scheme with a binder length range of 80–120 residues. Quercetin was processed under both partial-exposure and buried-ligand schemes, with binder length ranges of 90–140 and 100–150 residues, respectively.

For each design task, 100 candidate backbones were generated. The initial design cohort therefore consisted of 100 capsaicin-conditioned backbones, 100 (4R)- limonene-conditioned backbones, 100 partial-exposure quercetin-conditioned backbones and 100 buried quercetin-conditioned backbones.

RFdiffusion3 JSON metadata were used for initial structural filtering. Extracted or calculated metrics included raw chain breaks, expected chain breaks, additional chain breaks, backbone clashes, side-chain clashes, ligand-associated steric conflicts, minimum protein–ligand distance, maximum Cα deviation, radius of gyration, loop fraction, non-loop fraction, helix fraction, sheet fraction, number of secondary-structure elements, alanine content, glycine content and total residue number. Candidates with additional chain breaks or severe structural defects were excluded from ligand-centred pocket evaluation. Candidates with moderate side-chain clashes were retained only when the defect did not directly affect the ligand-proximal region.

### 2.4 Ligand-centred local pocket evaluation

JSON-passed candidates were evaluated by ligand-centred pocket analysis in PyMOL. This step assessed whether the generated backbone formed a physically plausible pocket or groove around the ligand and whether the pocket matched the intended exposure-conditioning scheme.

The quantitative descriptors used for this evaluation included buried ligand atom contact count, exposed ligand atom contact count where applicable, number of protein residues within 4.0 and 5.0 Å of the ligand, and severe ligand–protein clash count within 1.8 Å. Candidates were retained for sequence design when they showed sufficient contact with predefined buried atoms, appropriate accessibility of exposed or partially exposed atoms, a defined local pocket or groove, and no severe ligand-associated steric clash. Candidates were excluded or deprioritized when they showed insufficient buried- region contact, no clear local pocket, excessive burial of exposed atoms, excessive exposure of buried atoms or severe ligand–protein overlap.

For (4R)-limonene, pocket evaluation emphasized compact hydrophobic enclosure because the ligand lacked conventional hydrogen-bond donors or acceptors. For capsaicin and partial-exposure quercetin designs, evaluation considered both hydrophobic enclosure and retention of polar-atom accessibility. For buried quercetin designs, evaluation emphasized scaffold enclosure and absence of severe steric conflict.

### 2.5 LigandMPNN sequence design and RoseTTAFold3 structural back-prediction

Amino acid sequences were designed for all pocket-qualified backbones using LigandMPNN [14]. The RFdiffusion3-derived backbone geometry and ligand coordinates were used as the fixed structural context, and amino acid identities in the designed protein chain were optimized in the presence of the ligand-associated non-protein atomic environment. Ten sequences were generated for each retained backbone.

Each LigandMPNN-designed sequence was then submitted for RoseTTAFold3-based structural back-prediction [15]. These predictions were used to evaluate sequence foldability and structural consistency. Because the back-predicted structures did not contain ligand coordinates, this step assessed apo structural preservation rather than direct holo-complex prediction.

For each candidate, the RoseTTAFold3-predicted protein structure was aligned to the corresponding RFdiffusion3 reference backbone using matched Cα atoms. The same global Cα-based transformation was used for all downstream pocket comparisons. Global Cα RMSD and global backbone RMSD were calculated to assess overall fold preservation. The reference pocket was defined as all residues with at least one heavy atom within 6.0 Å of any ligand heavy atom in the RFdiffusion3 reference complex. Pocket Cα RMSD and pocket backbone RMSD were then calculated using these reference-defined pocket residues.

### 2.6 Projected ligand-pose compatibility and contact recovery analysis

Because RoseTTAFold3 back-predicted structures lacked ligand coordinates, the original RFdiffusion3 ligand pose was projected into the aligned coordinate frame of each predicted protein. Projected compatibility was assessed by comparing the aligned predicted protein with the original ligand coordinates.

The minimum projected protein–ligand heavy-atom distance was calculated for each candidate. A severe projected ligand clash was defined as any protein–ligand heavy-atom pair separated by less than 2.0 Å. Backbone–ligand and side-chain–ligand clash counts were recorded separately, with backbone atoms defined as N, Cα, C and O atoms. The total projected clash count was calculated as the sum of backbone–ligand and side-chain–ligand clashes.

Reference ligand-contact residues were defined as residues with at least one heavy atom within 4.5 Å of any ligand heavy atom in the RFdiffusion3 reference complex. Projected ligand-contact residues were defined using the same 4.5 Å cutoff after projection of the reference ligand pose into the aligned RoseTTAFold3-predicted protein structure. The projected contact recovery rate was calculated as:

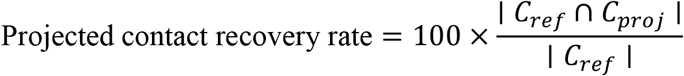

where *C_ref_*represented the reference ligand-contact residue set and *C_propp_*represented the projected ligand-contact residue set. The projected new contact rate was calculated as:

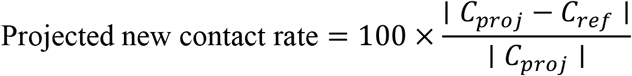

When *C_propp_*was empty, the projected new contact rate was not calculated. Carbon– carbon proximity pairs within 4.0 Å were also recorded as an approximate descriptor of nonpolar packing, rather than as a direct energetic measure of hydrophobic interaction.

### 2.7 Sequence-level property assessment and representative candidate selection

Sequence-level properties were calculated for LigandMPNN-designed candidates. NetSolP-1.0 with the ESM1b-based prediction model was used to estimate solubility [21] and usability for soluble expression and purification in *Escherichia coli*. ExPASy ProtParam was used to calculate the instability index and estimated *E. coli* half-life [22].

Integrated selection was performed separately within each parental backbone group because each retained backbone generated 10 designed sequences and 10 back-predicted structures. Candidate information from NetSolP, ProtParam and structural compatibility analysis was merged using candidate identifiers.

Candidates were classified into three decision categories. A candidate was classified as Strict selected when it was predicted to be stable, showed no projected ligand clash and achieved 100% projected contact recovery. When multiple strict candidates were available for the same backbone, they were ranked by higher ligand-interface sequence recovery, lower pocket backbone RMSD, lower instability index, higher NetSolP usability, higher NetSolP solubility and higher overall sequence recovery.

When no strict candidate was available, a candidate was classified as Conditional selected if it was predicted to be stable and showed no projected ligand clash. Conditional candidates were ranked by higher projected contact recovery, higher ligand-interface sequence recovery, lower pocket backbone RMSD, lower instability index, higher NetSolP usability, higher NetSolP solubility and higher overall sequence recovery. Conditional candidates were flagged when projected contact recovery was below 90% or pocket backbone RMSD exceeded 1.0 Å.

Candidates that did not satisfy the Strict or Conditional criteria were assigned to a repair-oriented or rejected category. Repair-oriented candidates were recorded when the defect could plausibly be addressed by local redesign or redocking. Rejected candidates were not included in the principal docking set. One representative candidate was retained from each parental backbone group

### 2.8 Molecular docking and docking-pose evaluation

Selected representative candidates were docked with their corresponding ligands using AutoDock Vina 1.2.7 [17]. RoseTTAFold3-selected protein structures were used as rigid receptors. Receptors were prepared in PDBQT format using Meeko 0.7.1, and ligand PDBQT files were generated from the prepared ligand structures while preserving ligand identity and chemical connectivity.

For each receptor, the docking box was centred on the corresponding mapped RFdiffusion3 reference ligand position. The docking box size was set to 20 × 20 × 20 Å. Docking was performed using the Vina scoring function with an exhaustiveness value of 32, a maximum of 10 output poses and an energy range of 4 kcal/mol.

Docking poses were compared with the corresponding RFdiffusion3 reference ligand pose mapped into the receptor coordinate frame. Docking-pose evaluation recorded the Vina score, reference-pose RMSD, recovery of reference protein–ligand contacts, formation of new contact residues and severe protein–ligand clashes. The prioritized docking pose was interpreted using design-consistency criteria rather than Vina score alone. Docking scores were used only for computational prioritization and were not interpreted as experimentally validated binding affinities.

Strict and Conditional representatives constituted the principal docking set. Repair- oriented candidates were not considered priority candidates but were subjected to exploratory redocking when ligand repositioning could potentially relieve projected local incompatibility.

### 2.9 Molecular dynamics trajectory analysis

To further evaluate the dynamic stability of selected capsaicin-binder complexes, molecular dynamics simulations were performed using GROMACS [23]. For each ligand–binder complex, the docked complex structure was used as the starting model, followed by system preparation, solvation, ion addition, energy minimization, equilibration, and production simulation under periodic boundary conditions. Each complex was independently simulated twice, with each replicate run lasting 100 ns, to assess the reproducibility and stability of the predicted binding mode. The resulting trajectories were analyzed to evaluate both the structural stability of the designed binder and the positional stability of the ligand within the binding pocket. Protein backbone RMSD was calculated to assess the overall conformational stability of the binder, whereas ligand heavy-atom RMSD was calculated after protein-backbone alignment to determine whether the small molecule remained stably positioned relative to the designed binding pocket. In addition, ligand–protein minimum distance and pocket- residue contact occupancy were analyzed to further characterize ligand retention and dynamic interaction persistence during the simulations. All trajectory-derived metrics were plotted as time-dependent profiles, and moving-average smoothing was applied where appropriate to facilitate visualization of overall trends while retaining the original trajectory-derived data for quantitative interpretation.

## 3 Results

### 3.1 Local pocket evaluation identified 31 capsaicin-binding backbone candidates meeting the screening criteria

A total of 100 candidate capsaicin-binding backbones were initially subjected to JSON- based structural quality assessment. Quality classification of all generated candidates showed that 68 models were graded as Good and 12 models as Acceptable, whereas 11 models were categorized as Poor extra chainbreak because of additional chain discontinuities and 9 models were categorized as Medium sidechain clash because of moderate side-chain steric conflicts. After excluding the 11 models with additional chain discontinuities and three models with ligand-proximal side-chain clashes, 86 models were retained for ligand-centred PyMOL-based local pocket evaluation. These results indicate that a substantial proportion of the generated candidate backbones exhibited acceptable structural continuity and lacked major geometric defects, allowing further assessment of their local binding environments around capsaicin.

The 86 models retained after JSON-based screening were subsequently evaluated by PyMOL-based local pocket evaluation (Figure 1, Table 3). None of the examined candidates exhibited severe ligand-related steric clashes within 1.8 Å, indicating that these structures generally met the basic requirement for spatial compatibility. Nevertheless, marked differences were observed among the candidate backbones in their contacts with the predefined buried region of capsaicin, the status of the exposed region, and their ability to form a local binding pocket. Following comprehensive evaluation, 31 models were classified as Good pocket, 25 models as Acceptable manual check, 23 models as Fail low buried contact because of insufficient contacts with the buried ligand atoms, and 1 model as Fail no clear pocket because no distinct local pocket was formed. In addition, 3 models were labelled Warning exposed overburied because the predefined exposed region may have been excessively covered, whereas another 3 models were labelled Warning exposed too free because of insufficient local support around the exposed region.

**Figure 1.**
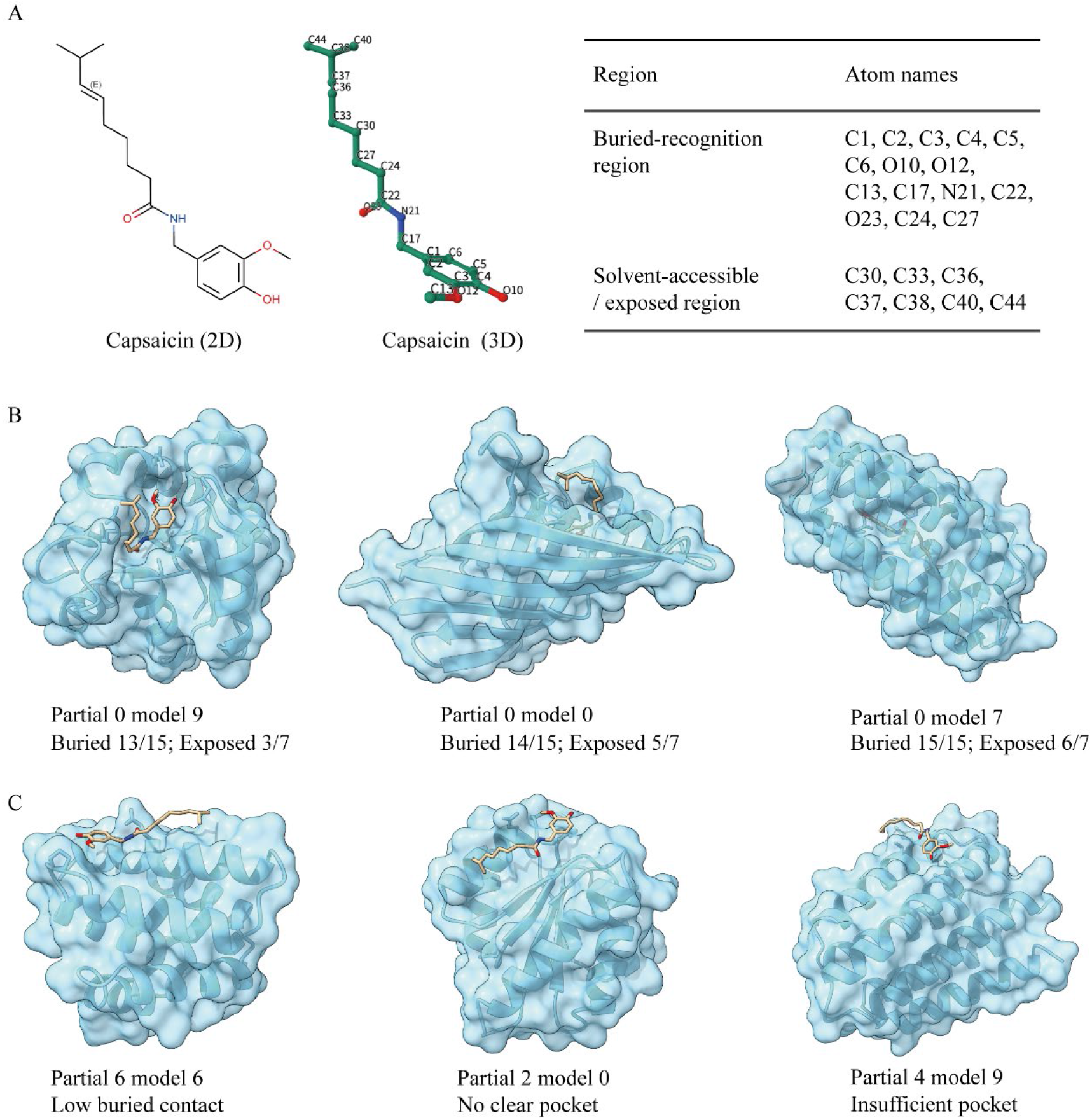
Partial-exposure-guided screening of candidate capsaicin-binding protein backbones. Note: (A) Two- and three-dimensional representations of capsaicin and the predefined buried-recognition and solvent-accessible regions used for ligand conditioning. (B) Representative selected backbones forming suitable local pockets around capsaicin. (C) Representative non-selected backbones with low buried-region contact, no clear pocket, or insufficient pocket formation. Protein backbones are shown in cyan, and capsaicin is shown as sticks.

**Table 3.** Top 10 candidate capsaicin-binding backbones after refined PyMOL-based screening.

| Candidate | Score | Buried | Exposed | Pocket 4 | Pocket 5 | Clash1.8 | Evaluation |
| --- | --- | --- | --- | --- | --- | --- | --- |
| partial 5 model 4 | 16.5538 | 13/15 | 4/7 | 16 | 22 | 0 | Good pocket |
| partial 0 model 7 | 17.2661 | 15/15 | 6/7 | 15 | 25 | 0 | Good pocket |
| partial 4 model 6 | 24.9440 | 13/15 | 5/7 | 15 | 22 | 0 | Good pocket |
| partial 0 model 9 | 25.1374 | 13/15 | 3/7 | 11 | 16 | 0 | Good pocket |
| partial 1 model 8 | 26.0098 | 14/15 | 2/7 | 13 | 20 | 0 | Good pocket |
| partial 3 model 1 | 26.7488 | 13/15 | 3/7 | 12 | 15 | 0 | Good pocket |
| partial 7 model 0 | 27.1878 | 14/15 | 6/7 | 12 | 17 | 0 | Good pocket |
| partial 7 model 7 | 27.5888 | 13/15 | 3/7 | 11 | 20 | 0 | Good pocket |
| partial 6 model 2 | 30.7796 | 12/15 | 4/7 | 11 | 20 | 0 | Good pocket |
| partial 1 model 2 | 31.4546 | 14/15 | 5/7 | 12 | 17 | 0 | Good pocket |

Note: Buried represents the number of buried atoms of capsaicin that were predefined to form contacts with the protein backbone, whereas exposed represents the number of atoms in the predefined solvent-exposed region that form contacts with the protein backbone. Because a partial-design strategy was adopted in this study, the exposed region was expected to retain a certain degree of solvent accessibility; therefore, candidates with either excessively high or excessively low exposed contact values required further manual inspection. Pocket 4 indicates the number of pocket residues surrounding capsaicin within 4 Å, and clash1.8 indicates the number of severe atomic clashes within 1.8 Å.

All 31 models classified as Good pocket were free of severe ligand-related steric clashes. Among them, the top 20 candidates contacted 12 to 15 predefined buried atoms, contained 8 to 16 pocket residues within 4 Å of capsaicin, and contained 13 to 25 pocket residues within 5 Å of the ligand. These results show that the selected candidates formed relatively continuous local environments around capsaicin and met the screening requirements for the partial burial design strategy. Among the candidates ranked by PyMOL-based local pocket evaluation, capsaicin partial 5 model 4 achieved the highest rank, with a final composite score of 16.5538. This model contacted 13/15 predefined buried atoms and 4/7 predefined exposed atoms, contained 16 and 22 pocket residues within 4 Å and 5 Å of capsaicin, respectively, and showed no severe steric clashes. Capsaicin partial 0 model 7, ranked second, contacted all 15/15 predefined buried atoms and contained 15 and 25 pocket residues within 4 Å and 5 Å of the ligand, respectively. The third- to fifth-ranked candidates, capsaicin partial 4 model 6, capsaicin partial 0 model 9, and Capsaicin partial 1 model 8, were also classified as Good pocket and contacted 13/15, 13/15, and 14/15 predefined buried atoms, respectively.

Initial structural quality scoring identifies candidate backbones with generally acceptable geometry, whereas ligand-centred PyMOL-based local pocket evaluation further distinguishes models that form capsaicin-binding environments meeting the predefined screening criteria. For example, capsaicin partial 4 model 9, which ranked first during the initial JSON-based screening because of its lowest initial composite score, contained only 7 and 10 pocket residues within 4 Å and 5 Å of capsaicin, respectively, and was therefore classified as Acceptable manual check rather than as a top pocket-forming candidate. Similarly, capsaicin partial 6 model 6, which ranked second in the initial screening, contacted only 7/15 predefined buried atoms and was consequently classified as Fail low buried contact.

Overall, 31 of the 100 initially generated capsaicin-binding backbones were classified as Good pocket after sequential initial quality screening and PyMOL-based local pocket evaluation. These candidates exhibited acceptable basic geometry, lacked severe ligand-related steric clashes, and formed local pockets around capsaicin that met the predefined screening criteria. Therefore, the 31 Good pocket backbones were selected for subsequent sequence design and downstream structural evaluation.

### 3.2 Local pocket evaluation identified 75 (4R)-limonene-binding backbone candidates meeting the screening criteria

A JSON-based structural quality assessment was performed for the 100 candidate (4R)- limonene-binding backbones generated. The results showed that 78 models were classified as Good, 12 as Acceptable, 5 as Poor extra chainbreak because of additional chain discontinuities, and another 5 as Medium sidechain clash because of moderate side-chain steric conflicts. Overall, 90% of the candidate backbones were rated as Good or Acceptable, indicating that RFdiffusion3 generated a high proportion of candidates with acceptable baseline structural quality under the buried-design setting for (4R)- limonene.

In the JSON-based composite-score ranking, (4R)-limonene buried 2 model 6 ranked first, with a composite score of 11.4022, a minimum ligand distance of 3.71 Å, a maximum Cα deviation of 0.055 Å, and a radius of gyration of 13.43 Å. This model was predominantly composed of α-helical secondary structure, suggesting favorable overall backbone regularity. The second- to fourth-ranked models, namely (4R)- limonene buried 5 model 6, (4R)-limonene buried 7 model 0, and (4R)-limonene buried 4 model 9, also showed low composite scores, no obvious major structural clashes, and high proportions of regular secondary structure, supporting their inclusion in subsequent ligand-centered PyMOL-based local pocket evaluation.

Following JSON prescreening, 95 candidate backbones were subjected to PyMOL-based local pocket evaluation. At this stage, only the five models with additional chain discontinuities were excluded, whereas models with moderate side-chain clashes were retained for visual inspection to determine whether these clashes affected the ligand-binding region. Because (4R)-limonene was designed under a buried-ligand constraint, this refinement focused on whether the predefined buried ligand atoms were sufficiently contacted by the candidate backbone and whether a continuous, clash-free hydrophobic pocket was formed around the ligand. None of the models subjected to PyMOL evaluation exhibited severe ligand–protein atomic clashes within 1.8 Å, indicating that the pre-screened candidates generally possessed acceptable basic spatial compatibility. Further classification showed that 61 models were rated as Excellent hydrophobic pocket, 14 as Good hydrophobic pocket, 16 as Acceptable manual check, and 4 as Fail low buried contact because of insufficient contacts with the predefined buried ligand atoms.

Notably, PyMOL-based local pocket evaluation changed the ranking of the leading candidates (Figure 2, Table 4). The highest-ranked model after JSON pre-screening, (4R)-limonene buried 2 model 6, contacted only 6/10 predefined buried atoms in the PyMOL analysis and contained 8 and 12 pocket residues within 4 Å and 5 Å of the ligand, respectively. Consequently, this model was classified as Acceptable manual check and did not remain among the optimal hydrophobic-pocket candidates. Similarly, (4R)-limonene buried 7 model 0, which ranked third in the JSON-based analysis, contacted only 7/10 predefined buried atoms and was also classified as Acceptable manual check.

**Figure 2.**
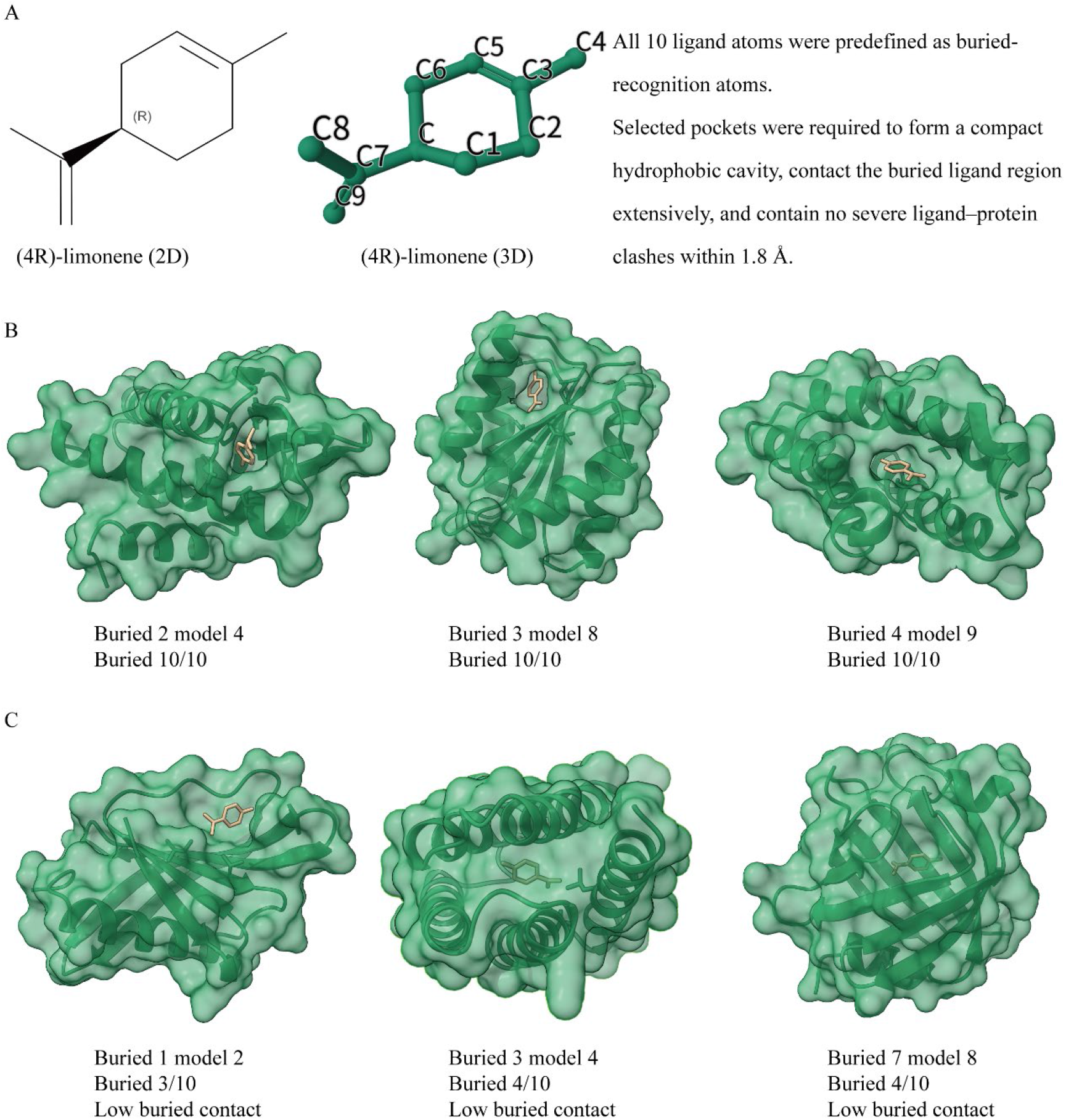
Buried-ligand-guided screening of candidate (4R)-limonene-binding protein backbones. Note: (A) Two- and three-dimensional structures of (4R)-limonene and the buried-ligand design strategy. (B) Representative selected backbones forming compact hydrophobic pockets around (4R)-limonene. (C) Representative non-selected backbones with insufficient contact with the predefined buried ligand region. Protein structures are shown in green, and (4R)-limonene is shown as sticks.

**Table 4.** Top 10 candidate (4R)-limonene-binding backbones after refined PyMOL-based screening.

| Candidate | Score | Buried | Pocket 4 | Pocket 5 | Clash 1.8 | Evaluation |
| --- | --- | --- | --- | --- | --- | --- |
| buried 5 model 6 | 12.5250 | 10/10 | 13 | 14 | 0 | Excellent hydrophobic pocket |
| buried 4 model 9 | 13.4089 | 10/10 | 11 | 14 | 0 | Excellent hydrophobic pocket |
| buried 6 model 3 | 19.6996 | 10/10 | 10 | 11 | 0 | Excellent hydrophobic pocket |
| buried 3 model 8 | 21.0215 | 10/10 | 10 | 13 | 0 | Excellent hydrophobic pocket |
| buried 2 model 4 | 22.8894 | 10/10 | 9 | 13 | 0 | Excellent hydrophobic pocket |
| buried 9 model 4 | 23.1448 | 10/10 | 9 | 14 | 0 | Excellent hydrophobic pocket |
| buried 8 model 4 | 23.2216 | 10/10 | 7 | 15 | 0 | Excellent hydrophobic pocket |
| buried 6 model 5 | 24.3867 | 10/10 | 8 | 13 | 0 | Excellent hydrophobic pocket |
| buried 5 model 1 | 24.4771 | 10/10 | 11 | 13 | 0 | Excellent hydrophobic pocket |
| buried 8 model 7 | 26.1713 | 10/10 | 9 | 15 | 0 | Excellent hydrophobic pocket |

Note: Contacted buried atoms indicate the number of predefined buried atoms of (4R)-limonene that formed local contacts with chain A in the designed backbone. Pocket residues within 4 Å and 5 Å represent the numbers of chain A residues located within the corresponding distance thresholds from the ligand. Severe clashes indicate the number of ligand–protein atomic contacts within 1.8 Å. Because (4R)-limonene was designed using a buried-ligand strategy, the completeness of buried-atom contacts and the formation of a continuous hydrophobic pocket were considered the primary criteria for candidate selection.

In contrast, (4R)-limonene buried 5 model 6, which ranked second in the JSON-based analysis, emerged as the highest-ranked candidate after PyMOL refinement. This model achieved a final score of 12.5250, contacted all 10/10 predefined buried atoms, contained 13 and 14 pocket residues within 4 Å and 5 Å of the ligand, respectively, and showed no severe atomic clashes. It was therefore classified as Excellent hydrophobic pocket. The model (4R)-limonene buried 4 model 9, initially ranked fourth by JSON analysis, also contacted all 10/10 buried atoms, contained 11 and 14 pocket residues within 4 Å and 5 Å, respectively, and achieved a final score of 13.4089, ranking second after PyMOL refinement. In addition, the third- to sixth-ranked models after refinement, (4R)-limonene buried 6 model 3, (4R)-limonene buried 3 model 8, (4R)-limonene buried 2 model 4, and (4R)-limonene buried 9 model 4, all achieved complete contact with the predefined buried atoms and formed local hydrophobic pockets without detectable severe steric clashes.

Overall, among the 100 initially generated candidate backbones for (4R)-limonene, 75 models passed the local-pocket evaluation, including 61 Excellent hydrophobic pocket models and 14 Good hydrophobic pocket models. These candidates showed no severe spatial clashes and formed reasonable hydrophobic pockets around (4R)-limonene to varying degrees.

### 3.3 Local pocket evaluation identified 56 quercetin-binding backbone candidates meeting the screening criteria

We performed an initial JSON-based structural quality assessment of the 200 candidate quercetin-binding backbones generated. Among all candidate models, 108 were classified as Good, 46 as Acceptable, 36 as Poor extra chainbreak because of additional chain breaks, and 10 as Medium sidechain clash because of moderate side-chain clashes.

All of the top 10 candidates ranked by the JSON-based composite score were classified as Good, and none showed raw chain breaks, additional chain breaks, backbone clashes, side-chain clashes, or ligand clashes. Among them, quercetin partial 5 model 7 ranked first in the JSON prescreening stage, with a composite score of 12.9316, a minimum ligand distance of 5.04 Å, a maximum Cα deviation of 0.062 Å, and a radius of gyration of 14.88 Å; this model was predominantly composed of α-helical structure. The models ranked second to fifth, namely quercetin partial 3 model 8, quercetin buried 8 model 1, quercetin partial 0 model 7, and quercetin buried 0 model 5, also exhibited relatively low composite scores and favorable overall structural quality. Because the quercetin candidate backbones comprised both partial and buried design modes, which impose distinct requirements on the local spatial environment of the ligand, PyMOL was subsequently used to evaluate their actual pocket-forming ability.

Following JSON prescreening, 154 candidate backbones were subjected to PyMOL-based local pocket evaluation, including 79 partial models and 75 buried models. PyMOL analysis revealed marked differences between the two design modes in their ability to form local quercetin-binding pockets. Among the 154 evaluated models, 33 were classified as Excellent quercetin buried pocket, 22 as Good quercetin buried pocket, and 1 as Good quercetin partial pocket. Therefore, a total of 56 candidate backbones passed the PyMOL-based local pocket evaluation. Among the remaining models, 17 partial models and 11 buried models required further manual inspection; 58 models were classified as Fail low buried contact because of insufficient contacts with buried atoms; 2 models were classified as Fail no clear pocket because they failed to form a clearly defined pocket; 4 models were classified as Fail severe clash because of severe steric clashes; and 6 partial models were classified as Warning partial exposed overburied because their predefined exposed regions were excessively buried.

A direct comparison of the two design strategies showed that the quercetin buried models had a substantially higher refinement pass rate. Among the 75 buried candidates, 55 were classified as either Excellent quercetin buried pocket or Good quercetin buried pocket, corresponding to a pass rate of 73.3%. By contrast, among the 79 partial candidates, only quercetin partial 3 model 8 was classified as Good quercetin partial pocket, corresponding to a pass rate of only 1.3%. These results indicate that, under the design constraints and screening criteria used in this study, the buried strategy was more suitable than the partial strategy for generating backbones capable of forming continuous local binding pockets around quercetin. In contrast, models generated using the partial strategy were more prone to insufficient contacts with buried regions or excessive burial of predefined exposed regions.

The final PyMOL-based composite ranking further supported the advantage of the buried design strategy (Figure 3, Table 5). Among the top 20 models in the final ranking, 19 originated from the quercetin buried design task, whereas only 1 originated from the quercetin partial design task. Of these top 20 models, 15 were classified as Excellent quercetin buried pocket, 4 as Good quercetin buried pocket, and 1 as Good quercetin partial pocket. For the buried models within the top 20, the number of contacted quercetin buried atoms ranged from 19/22 to 22/22, with an average contact proportion of approximately 92.3%. In addition, these models contained 10–16 pocket residues within 4 Å and 15–24 pocket residues within 5 Å of quercetin, indicating that they formed relatively continuous and compact local binding environments around the ligand.

**Figure 3.**
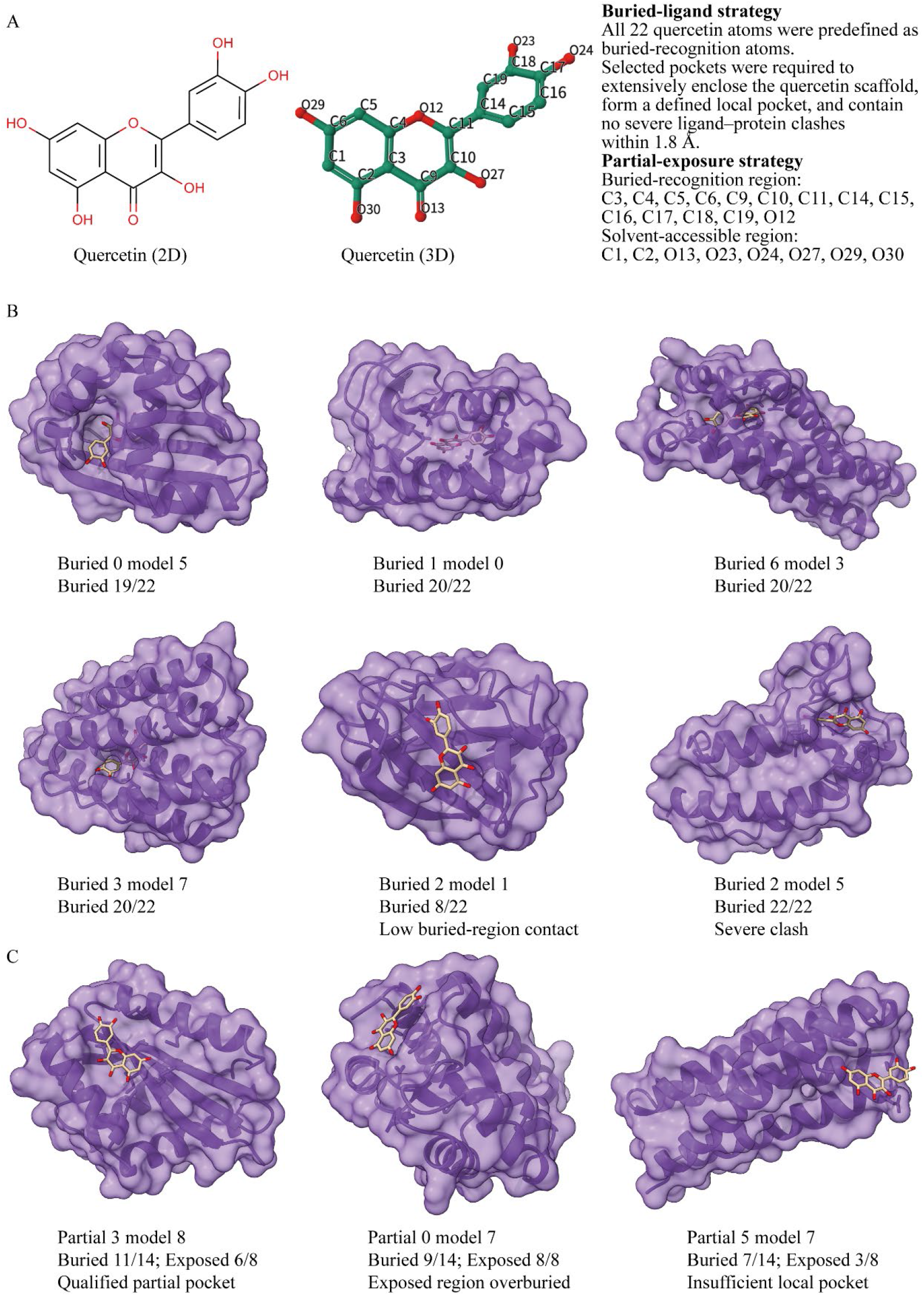
Comparison of buried-ligand and partial-exposure strategies for quercetin-binding backbone design. Note: (A) Quercetin structure and the buried-ligand and partial-exposure conditioning strategies. (B) Representative buried-mode backbones, including selected pockets with extensive quercetin enclosure and a non-selected structure with low buried-region contact. (C) Representative partial-mode backbones, including the qualified partial pocket and non-selected structures with excessive burial of the exposed region or insufficient local pocket formation. Protein structures are shown in purple, and quercetin is shown as sticks.

**Figure 4.**
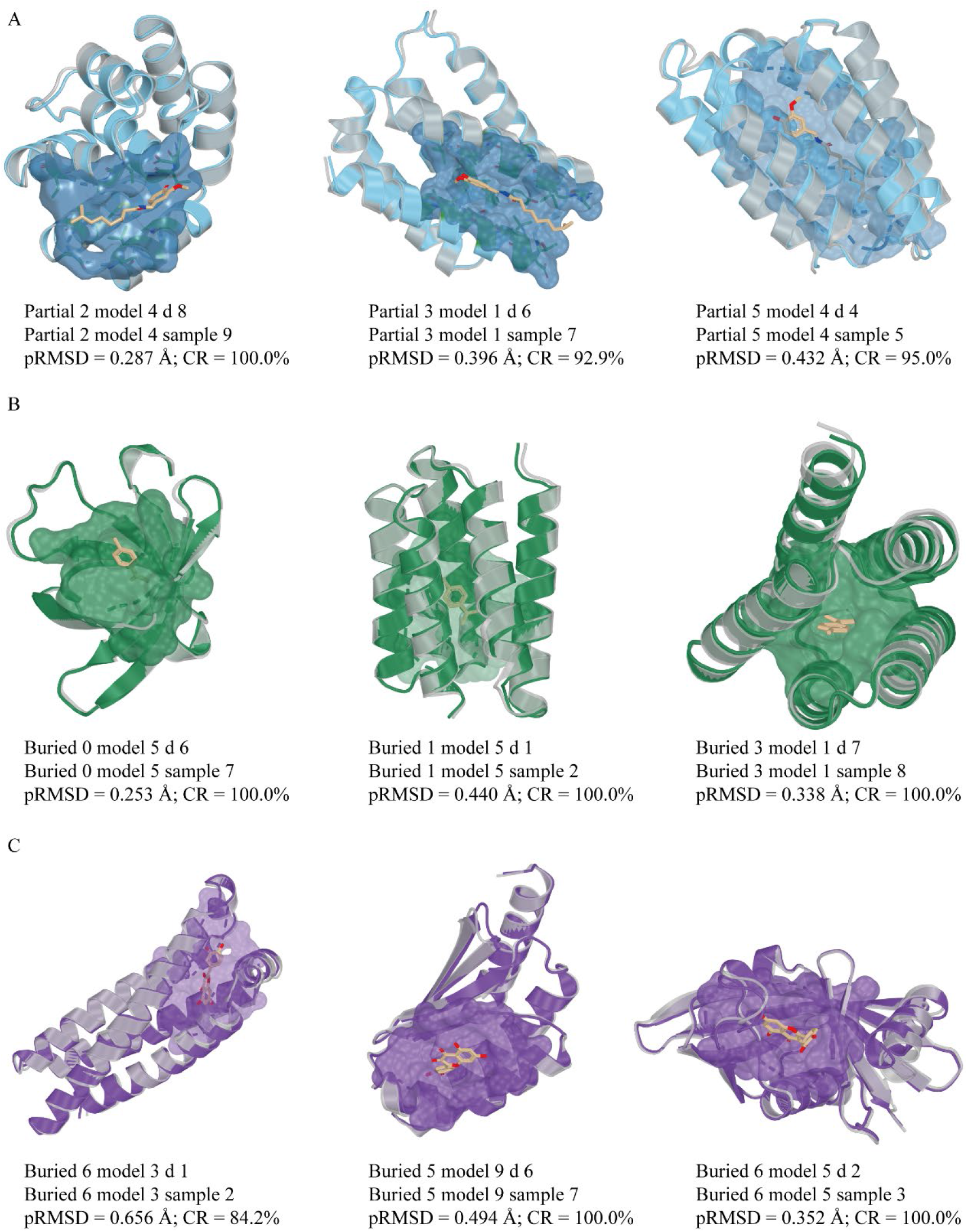
Structural preservation of representative ligand-binding pockets after sequence design and RoseTTAFold3 structural back-prediction. Note: Representative pocket overlays are shown for capsaicin (A), (4R)-limonene (B), and quercetin (C). For each ligand, three representative RoseTTAFold3 back-predicted candidate structures were superimposed onto their corresponding RFdiffusion3 reference designs to evaluate preservation of the designed local binding-pocket geometry. RoseTTAFold3 back-predicted protein structures are shown in blue, green, and purple for 4DY-, 9IR-, and QUE-binding candidates, respectively, whereas the corresponding RFdiffusion3 reference backbones are shown in grey. The original design-state ligand poses are shown as sticks, and ligand-contacting residues are displayed as sticks with semi-transparent local surfaces; non-contact regions are shown without surfaces. pRMSD indicates the pocket backbone RMSD between the RoseTTAFold3 back-predicted structure and its corresponding RFdiffusion3 reference design after structural superposition, and CR indicates recovery of the original ligand- contacting positions. All displayed candidates had a predicted ligand clash count of 0.

**Table 5.** Top 10 candidate quercetin-binding backbones after refined PyMOL-based screening.

| Candidate | score | Buried | Exposed | Pocket 4 | Pocket 5 | Clash 1.8 | Evaluation |
| --- | --- | --- | --- | --- | --- | --- | --- |
| buried 0 model 5 | 20.8095 | 19/22 | N/A | 10 | 15 | 0 | Good pocket |
| buried 5 model 8 | 22.2308 | 19/22 | N/A | 15 | 20 | 0 | Good pocket |
| buried 1 model 0 | 26.6031 | 20/22 | N/A | 13 | 23 | 0 | Excellent pocket |
| buried 6 model 3 | 27.0067 | 20/22 | N/A | 14 | 20 | 0 | Excellent pocket |
| buried 3 model 7 | 27.6116 | 20/22 | N/A | 14 | 20 | 0 | Excellent pocket |
| buried 6 model 7 | 27.6223 | 19/22 | N/A | 13 | 22 | 0 | Good pocket |
| buried 4 model 3 | 27.8524 | 22/22 | N/A | 15 | 22 | 0 | Excellent pocket |
| buried 4 model 4 | 28.7519 | 22/22 | N/A | 13 | 19 | 0 | Excellent pocket |
| buried 2 model 3 | 29.9434 | 20/22 | N/A | 13 | 17 | 0 | Excellent pocket |
| buried 7 model 7 | 30.2475 | 20/22 | N/A | 16 | 24 | 0 | Excellent pocket |

Note: For the buried design mode, all 22 quercetin ligand atoms were predefined as expected buried atoms; therefore, exposed atom contacts were not separately evaluated and are indicated as N/A. Buried atom contacts represents the number of predefined quercetin buried atoms forming local contacts with chain A. Pocket residues within 4 Å and Pocket residues within 5 Å indicate the numbers of chain A residues located within 4 Å and 5 Å of quercetin, respectively. *Severe clashes* refers to the number of ligand–protein atomic clashes detected within 1.8 Å. All top 10 candidates originated from the buried design mode, supporting its greater suitability for constructing candidate quercetin-binding pockets under the present screening criteria.

Following PyMOL-based local pocket evaluation, the ranking of candidate backbones changed markedly compared with the JSON prescreening results. Although quercetin partial 5 model 7 ranked first in the JSON prescreening stage and had the lowest initial composite score, it contacted only 7/14 predefined buried atoms and contained only 4 and 7 pocket residues within 4 Å and 5 Å of quercetin, respectively. Because it failed to form a sufficiently developed local binding environment around quercetin, this model was classified as Fail low buried contact. Similarly, quercetin partial 0 model 7, which ranked fourth in the JSON prescreening stage, contacted all 8/8 predefined exposed atoms, suggesting excessive burial of a region intended to retain partial solvent accessibility; accordingly, this model was classified as Warning partial exposed overburied. In contrast, models derived from the buried design task exhibited superior pocket-forming characteristics during PyMOL refinement. quercetin buried 0 model 5 ranked first after PyMOL refinement, with a final composite score of 20.8095. This model contacted 19/22 predefined buried atoms, contained 10 and 15 pocket residues within 4 Å and 5 Å of quercetin, respectively, and showed no severe steric clashes; it was therefore classified as Good quercetin buried pocket. The second-ranked model, quercetin buried 5 model 8, also contacted 19/22 buried atoms and formed local pockets containing 15 and 20 residues within 4 Å and 5 Å, respectively. The third- to fifth-ranked models, quercetin buried 1 model 0, quercetin buried 6 model 3, and quercetin buried 3 model 7, each contacted 20/22 predefined buried atoms and were classified as Excellent quercetin buried pocket, further demonstrating that the buried design strategy was more effective than the partial strategy in generating candidate backbones capable of extensively enclosing quercetin and forming continuous local binding pockets.

From the initial set of 200 quercetin candidate backbones, 56 models passed the PyMOL-based local pocket evaluation, including 33 Excellent quercetin buried pocket models, 22 Good quercetin buried pocket models, and 1 Good quercetin partial pocket model. Given the clear advantage of buried models in both screening pass rate and final composite ranking, the 55 buried candidates that passed refinement, together with the single qualified partial candidate, quercetin partial 3 model 8, will be used as inputs for subsequent LigandMPNN-based sequence design.

### 3.4 Ligand-dependent differences in local pocket formation

The local pocket evaluation results revealed clear ligand-dependent differences in the efficiency of generating candidate binding pockets. Among the three ligands, (4R)- limonene showed the highest local-pocket pass rate, with 75 of the 100 initially generated backbones passing PyMOL-based local pocket evaluation. This high success rate is consistent with the compact and highly hydrophobic nature of (4R)-limonene, as well as the buried-pocket design strategy used for this ligand. In contrast, capsaicin showed a lower pass rate, with 31 of 100 initially generated backbones classified as Good pocket. This likely reflects the structural complexity of capsaicin, which contains both a hydrophobic chain and polar functional groups, requiring a balance between hydrophobic enclosure and partial exposure of polar atoms.

Quercetin showed a more mode-dependent pattern. Although 56 of the 200 initially generated quercetin-binding backbones passed local pocket evaluation, the qualified candidates were highly enriched in the buried-design group. Among the 75 quercetin buried candidates subjected to PyMOL-based evaluation, 55 passed the local-pocket criteria, corresponding to a pass rate of 73.3%. By contrast, only 1 of the 79 quercetin partial candidates passed evaluation. These results suggest that the buried design strategy was more effective for quercetin under the screening criteria used in this study, whereas the partial-exposure strategy more frequently resulted in insufficient buried- region contacts or excessive burial of predefined exposed regions. Overall, these findings indicate that ligand size, hydrophobicity, rigidity, polarity, and the selected exposure-conditioning strategy jointly influence the efficiency of computational local pocket formation.

### 3.5 Integrated screening identifies candidate structures for docking analysis

For capsaicin, the 31 partial-pocket backbones retained from the preceding PyMOL-based local pocket evaluation produced 310 back-predicted structures. Multi-parameter screening retained one representative structure from each backbone, resulting in 31 candidate capsaicin-binding structures for subsequent molecular docking and further structural evaluation. All 31 representative candidates were predicted to be stable, with instability indices ranging from 10.56 to 39.98 and a median value of 29.74. Except for partial 3 model 8 d 0, for which the estimated Escherichia coli *in vivo* half-life was reported as unknown, the remaining 30 candidates had predicted half-lives of >10 h. NetSolP predictions showed that the retained candidates had solubility scores ranging from 0.5528 to 0.9824, with a median of 0.9213, and usability scores ranging from 0.4233 to 0.6982, with a median of 0.5237, suggesting generally favorable predicted solubility and potential expression usability.

Structural back-prediction analysis further indicated that most capsaicin candidates preserved the local geometry of the originally designed binding pocket. The pocket backbone RMSD values of the 31 representative structures ranged from 0.2872 to 3.1098 Å, with a median of 0.6624 Å; 24 candidates had pocket backbone RMSD values below 1.0 Å, and 29 candidates had values no greater than 2.0 Å. Evaluation of projected compatibility with the originally designed ligand pose showed that 25 candidates displayed no detectable projected protein–ligand steric clashes, accounting for 80.6% of all representative structures. Among these clash-free candidates, 16 also achieved a projected contact recovery rate of at least 90%, indicating that they retained both the local organization and the principal interaction pattern of the intended capsaicin-binding pocket after structural back-prediction.

The projected ligand-contact pattern was broadly maintained across the representative capsaicin candidates. Projected contact recovery rates ranged from 60.00% to 100.00%, with a median of 92.31%, and 17 candidates achieved contact recovery rates of at least 90%. In particular, partial 0 model 7 d 4, partial 2 model 3 d 6, partial 2 model 4 d 8, partial 2 model 6 d 8, and partial 8 model 2 d 8 showed no detectable ligand clashes and achieved 100% projected contact recovery. Among these candidates, partial 2 model 4 d 8 exhibited the lowest pocket backbone RMSD, at 0.2872 Å; partial 2 model 3 d 6 achieved the highest usability score, at 0.6982; and partial 8 model 2 d 8 combined 100% contact recovery with a pocket backbone RMSD of 0.4454 Å and a usability score of 0.6438, indicating favorable overall structural preservation and predicted usability.

Although representative candidate structures were obtained for all 31 capsaicin backbones, six candidates showed residual projected protein–ligand clashes. partial 4 model 7 d 9 and partial 8 model 6 d 5 each contained one residual clash, partial 3 model 8 d 0 contained two clashes, partial 1 model 2 d 6 and partial 1 model 7 d 1 each contained three clashes, and partial 1 model 8 d 2 showed the highest clash count, with 22 clashes. In addition, partial 3 model 8 d 0 and partial 8 model 6 d 5 exhibited relatively low contact recovery rates of 61.54% and 60.00%, respectively, indicating limited preservation of the original binding-pocket interaction pattern. All 31 representative capsaicin-binding structures were retained for docking-based reassessment, whereas the 25 clash-free candidates were considered the principal docking evaluation set. Of these, the 16 candidates simultaneously showing no clashes and high projected contact recovery were designated as the priority subset for subsequent evaluation. The six candidates with residual clashes require cautious interpretation in conjunction with subsequent redocking or local conformational optimization and were not considered priority structures for molecular dynamics simulation.

For (4R)-limonene, the 75 buried-pocket backbones retained after PyMOL-based local pocket evaluation generated 750 back-predicted structures. Following integrated screening, 28 backbones yielded strictly selected representative structures, accounting for 37.3% of the 75 local-pocket-passed backbones. All strictly selected candidates were predicted to be stable, showed no detectable projected protein–ligand clashes, and achieved 100% projected contact recovery. Their pocket backbone RMSD values ranged from 0.2526 to 0.9626 Å, with a median of 0.4848 Å, indicating that the predefined (4R)-limonene-binding pocket geometry was generally preserved after sequence design and structural back-prediction. The ligand-interface sequence recovery of these strictly selected structures ranged from 0.3636 to 0.8750, with a median of 0.6786, indicating that multiple candidate structures preserved local pocket geometry while also retaining sequence features similar to those of the original designed interface.

In addition to the strictly selected structures, 26 (4R)-limonene backbones yielded conditionally selected representative structures, corresponding to 34.7% of the 75 local- pocket-passed backbones. These candidates were also predicted to be stable and showed no detectable projected protein–ligand clashes, but they did not completely recover all projected ligand-contact positions. Their contact recovery rates ranged from 54.55% to 92.86%, with a median of 90.91%, whereas their pocket backbone RMSD values ranged from 0.3475 to 2.1772 Å, with a median of 0.4909 Å. Although most conditionally selected structures retained relatively small local pocket deviations, variation in contact recovery and pocket preservation remained evident. Therefore, these 26 structures were retained for subsequent docking-based screening and conformational reassessment, but were not directly prioritized for molecular dynamics simulation.

The remaining 21 (4R)-limonene backbones failed to yield representative structures that simultaneously satisfied the requirements of favorable predicted stability and absence of projected protein–ligand clashes, accounting for 28.0% of the 75 local- pocket-passed backbones. These backbones were classified as requiring repair or redesign and were not included in the current formal docking candidate set. Overall, 54 representative candidate (4R)-limonene-binding structures derived from 750 back-predicted structures were retained for subsequent molecular docking analysis, corresponding to 72.0% of the 75 local-pocket-passed backbones. Among them, the 28 strictly selected candidates, which simultaneously exhibited favorable predicted stability, absence of projected ligand clashes, complete projected contact recovery, and low pocket backbone deviation, were prioritized for subsequent molecular dynamics evaluation.

For quercetin, the 56 candidate backbones retained after PyMOL-based local pocket evaluation generated 560 back-predicted structures. Integrated screening identified 11 backbones yielding strictly selected representative structures. These candidates were predicted to be stable, displayed no detectable projected protein–ligand clashes with the originally designed quercetin ligand pose, and achieved 100% projected ligand-contact recovery, indicating effective preservation of the intended quercetin-binding pocket geometry. Their pocket-region backbone RMSD values ranged from 0.3515 to 0.8598 Å, with a median of 0.4987 Å, whereas their ligand-interface sequence recovery values ranged from 0.4118 to 0.6667, with a median of 0.5263. These strictly selected structures were therefore considered priority candidates for subsequent quercetin docking analysis.

A further 21 quercetin backbones yielded conditionally selected representative structures. These candidates were also predicted to be stable and exhibited no detectable projected protein–ligand clashes with the originally designed quercetin pose, but their contact recovery rates did not reach 100%, suggesting partial loss or rearrangement of original design contacts after structural back-prediction. Their pocket-region backbone RMSD values ranged from 0.3562 to 2.0665 Å, with a median of 0.5720 Å, whereas their projected ligand-contact recovery rates ranged from 75.0000% to 95.0000%, with a median of 88.8889%. Although these structures did not fully recover the intended quercetin-contact network, most retained relatively limited pocket deformation and acceptable spatial compatibility. Therefore, the 21 conditionally selected structures were retained together with the strictly selected structures for subsequent docking-based screening, followed by reassessment of the quercetin binding orientation, key interaction residues, and local pocket reorganization.

The remaining 24 quercetin backbones did not yield representative structures suitable for direct entry into molecular docking and were classified as requiring repair or redesign. Among these, all 10 back-predicted structures derived from partial 3 model 8 contained at least one predicted side-chain clash with quercetin and therefore failed to provide a directly usable representative structure for subsequent analysis. Within this group, partial 3 model 8 d 3 was retained as a starting structure for further repair. This candidate was predicted to be stable, had an estimated *E. coli* half-life of >10 h, and showed NetSolP solubility and usability scores of 0.9273 and 0.7147, respectively. Its pocket-region backbone RMSD was 0.4710 Å, its projected ligand-contact recovery rate was 81.8182%, and it contained only one side-chain–quercetin clash, representing the lowest clash burden among the stable candidates derived from this backbone. Although this structure was not directly advanced to docking or molecular dynamics simulation, its overall features support its use as a priority starting point for local side-chain optimization, redesign, or conformational refinement.

In total, one representative structure was retained from each of the 56 quercetin backbones, yielding a representative candidate set of 56 structures. Among them, 11 were classified as strictly selected and 21 as conditionally selected, resulting in 32 representative structures suitable for subsequent quercetin docking-based screening. The remaining 24 representative structures were excluded from the current docking and molecular dynamics workflow because of insufficient stability, residual ligand clashes, or failure to satisfy both pocket-preservation and basic quality-control criteria, and were instead retained as candidates for subsequent structural repair and redesign.

Overall, multi-parameter integrated screening identified distinct candidate sets for the three ligands (Table 6). For capsaicin, 31 representative structures were retained for docking-based reassessment, among which 25 were free of projected ligand clashes and 16 simultaneously showed no ligand clashes and high projected contact recovery. For (4R)-limonene, 54 representative structures were retained for subsequent docking-based screening, including 28 strictly selected candidates with predicted stability, absence of projected ligand clashes, complete projected contact recovery, and low pocket backbone deviation. For quercetin, 32 representative structures were retained for docking-based screening, including 11 strictly selected candidates satisfying the requirements of clash-free geometry and complete contact recovery. These prioritized candidate binder structures provide a structural basis for subsequent docking validation, conformational reassessment, and molecular dynamics-based evaluation of binding-pocket stability.

**Table 6.** Top 10 candidate structures for capsaicin, (4R)-limonene, and quercetin after LigandMPNN design and RoseTTAFold3 validation.

| Type | Candidate | Solubility | Usability | Instability index | RMSD | Contact recovery | New contact ratio |
| --- | --- | --- | --- | --- | --- | --- | --- |
| 4DY | partial 2 model 4 d8 | 0.9607 | 0.5567 | 29.20 | 0.2872 | 100.0000 | 8.3333 |
| 4DY | partial 3 model 1 d6 | 0.8724 | 0.4451 | 25.50 | 0.3957 | 92.8571 | 0.0000 |
| 4DY | partial 5 model 4 d4 | 0.9542 | 0.5166 | 37.35 | 0.4315 | 95.0000 | 17.3913 |
| 4DY | partial 8 model 2 d8 | 0.8742 | 0.6438 | 20.22 | 0.4454 | 100.0000 | 15.3846 |
| 4DY | partial 2 model 3 d6 | 0.9578 | 0.6982 | 10.56 | 0.4702 | 100.0000 | 5.5556 |
| 4DY | partial 0 model 7 d4 | 0.9399 | 0.4926 | 34.74 | 0.4847 | 100.0000 | 9.0909 |
| 4DY | partial 5 model 5 d7 | 0.8717 | 0.4950 | 39.98 | 0.5057 | 92.3077 | 25.0000 |
| 4DY | partial 9 model 5 d4 | 0.9824 | 0.6754 | 28.68 | 0.5224 | 92.3077 | 14.2857 |
| 4DY | partial 6 model 2 d8 | 0.9761 | 0.5237 | 33.41 | 0.5732 | 94.1176 | 20.0000 |
| 4DY | partial 2 model 6 d8 | 0.9659 | 0.5406 | 20.30 | 0.5770 | 100.0000 | 0.0000 |
| 9IR | buried 0 model 5 d6 | 0.9614 | 0.8252 | 25.39 | 0.2526 | 100.0000 | 0.0000 |
| 9IR | buried 5 model 2 d5 | 0.8676 | 0.3367 | 1.37 | 0.2594 | 100.0000 | 7.1429 |
| 9IR | buried 4 model 5 d4 | 0.9256 | 0.6361 | 27.03 | 0.3083 | 100.0000 | 23.0769 |
| 9IR | buried 3 model 1 d7 | 0.8863 | 0.5394 | 17.73 | 0.3375 | 100.0000 | 8.3333 |
| 9IR | buried 1 model 3 d0 | 0.7616 | 0.3933 | 37.40 | 0.3633 | 100.0000 | 9.0909 |
| 9IR | buried 8 model 9 d1 | 0.8796 | 0.4385 | 16.19 | 0.3674 | 100.0000 | 6.2500 |
| 9IR | buried 5 model 5 d4 | 0.9016 | 0.6253 | 23.09 | 0.3749 | 100.0000 | 15.3846 |
| 9IR | buried 2 model 8 d6 | 0.7992 | 0.4518 | 22.88 | 0.3858 | 100.0000 | 10.0000 |
| 9IR | buried 2 model 7 d4 | 0.8946 | 0.7007 | 33.13 | 0.4198 | 100.0000 | 16.6667 |
| 9IR | buried 2 model 3 d0 | 0.9117 | 0.5322 | 32.60 | 0.4359 | 100.0000 | 21.4286 |
| QUE | buried 6 model 5 d2 | 0.8010 | 0.3989 | 30.84 | 0.3515 | 100.0000 | 5.2632 |
| QUE | buried 1 model 0 d7 | 0.9140 | 0.7571 | 33.27 | 0.4451 | 100.0000 | 21.7391 |
| QUE | buried 4 model 0 d8 | 0.8908 | 0.5999 | 25.12 | 0.4776 | 100.0000 | 6.2500 |
| QUE | buried 5 model 9 d6 | 0.9168 | 0.6453 | 28.13 | 0.4942 | 100.0000 | 7.1429 |
| QUE | buried 0 model 5 d6 | 0.8091 | 0.4476 | 35.21 | 0.4979 | 100.0000 | 13.3333 |
| QUE | buried 5 model 6 d7 | 0.9322 | 0.6499 | 12.66 | 0.4987 | 100.0000 | 4.7619 |
| QUE | buried 7 model 7 d4 | 0.8922 | 0.3688 | 37.22 | 0.5188 | 100.0000 | 13.6364 |
| QUE | buried 7 model 9 d3 | 0.8286 | 0.4520 | 30.95 | 0.7848 | 100.0000 | 5.5556 |
| QUE | buried 8 model 1 d7 | 0.7214 | 0.3977 | 34.06 | 0.7950 | 100.0000 | 5.5556 |
| QUE | buried 5 model 1 d3 | 0.9093 | 0.5716 | 10.14 | 0.8004 | 100.0000 | 15.7895 |

Note: 4DY, 9IR, and QUE denote capsaicin-, (4R)-limonene-, and quercetin-binding candidates, respectively. Within each ligand group, candidates are ranked in ascending order of pocket backbone RMSD. Solubility and usability were predicted using NetSolP-1.0, whereas instability index and predicted intracellular half-life in *Escherichia coli* were estimated using ProtParam. The predicted half-life in *E. coli* was used as an auxiliary indicator of potential intracellular stability and expression feasibility in a prokaryotic expression system, rather than as a direct measure of experimental expression yield. Pocket backbone RMSD (Å) represents the deviation of backbone atoms within the ligand-binding pocket between each RoseTTAFold3 back-predicted structure and its corresponding RFdiffusion3 design after structural superposition. All candidate structures listed in this table had a ligand clash count of 0, indicating no detected steric conflict with the projected protein–ligand clashes. Contact recovery (%) represents the proportion of original ligand-contacting positions retained after RoseTTAFold3 structural back-prediction, whereas new contact ratio (%) represents the proportion of newly formed ligand-contacting positions relative to the predicted contact set.

### 3.6 Reference-pose-guided docking and pose recovery analysis

Among the 31 representative capsaicin-binding structures retained after integrated screening, 30 were successfully mapped to their corresponding reference binding pockets, whereas one candidate was excluded because a reliable structural correspondence could not be established owing to inconsistent Cα atom matching between the predicted structure and its reference backbone. For the 30 successfully mapped structures, the pocket-region RMSD ranged from 0.250 to 2.337 Å, with a median of 0.6165 Å, indicating that most candidates preserved the local geometry of the originally designed capsaicin-binding pocket.

Using a pocket-region RMSD threshold of <1.5 Å as a structural-preservation criterion for docking preparation, 27 capsaicin-binding structures were retained. Their pocket-region RMSD values ranged from 0.250 to 1.266 Å, with a median of 0.600 Å. The remaining three structures were excluded because of more pronounced local pocket deviations, with pocket RMSD values of 1.765, 1.994, and 2.337 Å, respectively. During receptor preparation, 19 of the 27 retained structures were successfully converted into standardized receptor models suitable for AutoDock Vina analysis. The remaining eight structures were excluded because abnormal residue–residue connections prevented reliable receptor parameterization without manual structural modification.

Focused docking of capsaicin into the 19 successfully prepared capsaicin-binding receptors generated valid docking poses for all candidates. The best-scoring poses showed predicted binding affinities ranging from −8.678 to −5.181 kcal mol⁻¹, with a median of −7.216 kcal mol⁻¹; 12 candidates achieved best-pose affinities of no higher than −7.0 kcal mol⁻¹. Based on affinity alone, partial 5 model 6 sample 1, partial 5 model 5 sample 8, partial 5 model 4 sample 5, partial 5 model 1 sample 10, and partial 7 model 4 sample 10 ranked highest, with predicted affinities of −8.678, −8.019, −7.942, −7.925, and −7.887 kcal mol⁻¹, respectively.

Among all capsaicin candidates, partial 5 model 4 sample 5 exhibited the most favorable integrated performance. Its selected mode-1 pose had a predicted affinity of −7.942 kcal mol⁻¹, an in-place heavy-atom RMSD of 1.0274 Å relative to the reference capsaicin pose, a reference-contact recovery rate of 88.89%, and a new contact proportion of only 15.79%, with no severe protein–ligand steric clashes detected. partial 5 model 1 sample 10 was also classified as a high-priority candidate, with a mode-1 affinity of −7.925 kcal mol⁻¹, an in-place RMSD of 1.1266 Å, a reference-contact recovery rate of 92.86%, and no severe steric clashes. Although this candidate showed slightly higher recovery of the original contacts than the leading candidate, its new contact proportion reached 27.78%, suggesting a greater degree of local interaction rearrangement.

Three additional structures, partial 2 model 4 sample 9, partial 0 model 9 sample 3, and partial 2 model 6 sample 9, were retained as supplementary candidates. partial 2 model 4 sample 9 recovered 90.00% of the reference contacts and showed an in-place RMSD of 1.4186 Å, but its affinity was relatively weak at −5.181 kcal mol⁻¹. partial 2 model 6 sample 9 likewise maintained a relatively high reference-contact recovery rate with a low proportion of newly formed contacts, although its predicted affinity was less favorable than those of the two high-priority candidates. By contrast, partial 0 model 9 sample 3 displayed a relatively favorable docking affinity but weaker restoration of the reference contact pattern and a higher proportion of new contacts, indicating a more pronounced alteration of the designed binding mode.

Notably, the candidate with the most favorable docking affinity was not the most suitable structure for downstream dynamic evaluation. partial 5 model 6 sample 1 achieved the lowest predicted affinity of −8.678 kcal mol⁻¹, but its clash-free representative pose still deviated substantially from the reference capsaicin position, with an in-place RMSD of 4.1796 Å and a new contact proportion of 56.25%. Similarly, partial 7 model 4 sample 10 achieved an affinity of −7.887 kcal mol⁻¹ but displayed an in-place RMSD of 8.3062 Å relative to the reference pose. These findings indicate that affinity-based ranking alone may favor alternative low-energy ligand orientations that deviate from the original design objective. Therefore, partial 5 model 4 sample 5 and partial 5 model 1 sample 10 were prioritized as the principal capsaicin-binding candidates for subsequent molecular dynamics analysis. Thus, the capsaicin candidate set was reduced from 31 representative structures to 19 formally docked receptors.

For (4R)-limonene, the 54 representative binder structures retained after integrated screening were subjected to reference-pocket-guided docking preparation. All 54 candidates successfully completed local pocket mapping and docking-box construction using the reference (4R)-limonene position, with a uniform docking region of 20 × 20 × 20 Å established around the reference ligand centre. During standardized receptor preparation, 48 candidate structures were successfully converted into receptor PDBQT files. Six candidates—buried 0 model 3 sample 4, buried 0 model 4 sample 8, buried 3 model 8 sample 6, buried 5 model 7 sample 8, buried 7 model 2 sample 1, and buried 8 model 6 sample 5—were excluded because abnormal residue–residue connections prevented reliable receptor preparation without manual intervention.

All 48 successfully prepared receptors completed AutoDock Vina docking and generated corresponding ligand-pose output files. Based on the mode-1 docking pose, buried 2 model 2 sample 8 achieved the most favorable predicted affinity, at −7.953 kcal mol⁻¹, followed by buried 8 model 3 sample 2 and buried 4 model 7 sample 10, with affinities of −7.829 and −7.775 kcal mol⁻¹, respectively. The top ten candidates ranked by affinity displayed acceptable local pocket alignment deviations, indicating that their favorable docking scores were not simply associated with major deformation of the reference pocket.

To determine whether the docked (4R)-limonene poses reproduced the intended design- state binding mode, the mode-1 poses were further evaluated using connectivity-aware in-place RMSD, ligand centre-of-mass displacement, reference-contact recovery and new contact formation. Of the 48 docked candidates, seven were classified as Tier A structures with strong support for reference-pose recovery, 13 were classified as Tier B structures with partial support, and 28 were classified as Tier C structures showing lower priority because of substantial pose or contact-network deviation.

Integration of docking affinity with pose and contact preservation. Among the Tier A candidates, buried 1 model 5 sample 2 displayed the most favorable overall balance of docking and design-pose recovery characteristics. Its predicted affinity was −6.964 kcal mol⁻¹, its connectivity-aware in-place RMSD was only 0.7057 Å, its ligand centre-of-mass displacement was 0.0495 Å, and its reference-contact recovery rate reached 93.3333%, with no newly formed contact residues detected. These results indicate that redocked (4R)-limonene returned closely to the intended reference position while largely preserving the original interaction pattern.

The candidate buried 9 model 9 sample 7 was also retained in the primary molecular dynamics shortlist. This candidate showed a predicted affinity of −6.813 kcal mol⁻¹, an in-place RMSD of 1.0878 Å, a centre-of-mass displacement of 0.7122 Å, and a contact recovery rate of 90.9091%, supporting its suitability as a parallel validation candidate. In contrast, buried 2 model 2 sample 8, although showing the most favorable predicted affinity of −7.953 kcal mol⁻¹ and a pocket-alignment RMSD of only 0.3185 Å, exhibited an in-place RMSD of 4.1806 Å and was therefore classified as Tier C.

Based on the combined assessment of docking affinity, reference-pose recovery, ligand displacement, and contact-network consistency, buried 1 model 5 sample 2 was selected as the primary (4R)-limonene candidate for subsequent molecular dynamics simulation. Its predicted protein structure together with the AutoDock Vina mode-1 pose (4R)- limonene pose was used to construct the starting protein–ligand complex for downstream dynamic evaluation.

For quercetin, 32 representative binder structures, including 11 strictly selected and 21 conditionally selected candidates, were retained after integrated screening and subjected to docking preparation. All 32 candidates were successfully aligned to their corresponding reference designs and used to generate local docking boxes centred on the design-state quercetin position. The aligned Cα RMSD values had a median of 0.6972 Å and a maximum value of 1.6741 Å, indicating that the overall conformations of most back-predicted structures remained close to their original designed backbones. Three candidates, buried 2 model 2 sample 0, buried 4 model 4 sample 6, and buried 5 model 3 sample 5, showed Cα RMSD values above 1.0 Å and were therefore considered candidates requiring particular attention during interpretation of their docked ligand poses.

During receptor PDBQT preparation, two quercetin candidates, buried 8 model 1 sample 8 and buried 8 model 3 sample 4, failed standardized processing because abnormal additional residue–residue connections were detected. These candidates were excluded from formal docking analysis to avoid introducing artificial pocket changes through manual structural correction. Consequently, 30 quercetin-binding receptors, including 10 strictly selected and 20 conditionally selected candidates, entered formal AutoDock Vina docking.

All 30 quercetin receptors successfully generated docking outputs, and their best-scoring ligand poses were retained for subsequent binding-mode assessment. The best predicted Vina affinities ranged from −9.284 to −5.279 kcal mol⁻¹, with a median of −7.965 kcal mol⁻¹. The connectivity-aware in-place RMSD values of the mode-1 quercetin poses ranged from 0.8355 to 13.5304 Å, with a median of 3.4933 Å, whereas ligand centre-of-mass displacements ranged from 0.2991 to 11.9480 Å, with a median of 1.5060 Å. The total design-state contact recovery rate ranged from 4.5455% to 95.2381%, with a median of 75.0000%. Among the 29 candidates containing comparable reference polar contacts, the polar-contact recovery rate ranged from 0% to 100%, with a median of 44.4444%. These results indicate substantial variation among quercetin candidates in their ability to reproduce the intended ligand-recognition mode following docking.

The candidate with the most favorable quercetin docking affinity did not exhibit the strongest reference-pose recovery. buried 6 model 5 sample 3 achieved the lowest predicted affinity among all formally docked candidates, at −9.284 kcal mol⁻¹; however, its mode-1 quercetin pose displayed an in-place RMSD of 3.4403 Å, a centre-of-mass displacement of 2.6163 Å, and a contact recovery rate of only 72.2222%, leading to its classification as Tier C. Similarly, buried 1 model 1 sample 1, buried 4 model 6 sample 7, and buried 3 model 6 sample 3 displayed predicted affinities more favorable than −8.97 kcal mol⁻¹ but showed substantial ligand-pose deviation or insufficient restoration of the design-state contact network. Thus, as observed for capsaicin and (4R)-limonene, affinity alone was insufficient to identify quercetin candidates that preserved the intended binding mode.

Integrated evaluation of affinity, quercetin pose recovery, centre-of-mass displacement, total contact recovery, and polar-contact recovery classified five candidates as Tier A, four as Tier B, and 21 as Tier C. The Tier A candidates were buried 6 model 3 sample 2, buried 1 model 0 sample 8, buried 4 model 3 sample 8, buried 6 model 1 sample 5, and buried 5 model 6 sample 8. Among them, buried 6 model 3 sample 2 displayed a favorable predicted affinity of −9.082 kcal mol⁻¹, a total contact recovery rate of 94.4444%, and complete restoration of reference polar contacts. buried 1 model 0 sample 8, which originated from the strictly selected group, showed an affinity of −8.635 kcal mol⁻¹, an in-place RMSD of 1.1457 Å, a centre-of-mass displacement of 0.4121 Å, a total contact recovery rate of 91.3043%, and a polar-contact recovery rate of 81.8182%. buried 4 model 3 sample 8 showed the lowest quercetin in-place RMSD, at 0.8355 Å, together with total and polar-contact recovery rates of 85.0000% and 75.0000%, respectively. buried 5 model 6 sample 8, also derived from the strictly selected group, showed an in-place RMSD of 0.8982 Å and a total contact recovery rate of 95.2381%. buried 6 model 1 sample 5 exhibited the smallest centre-of-mass displacement, at 0.2991 Å; however, because its reference design contained no comparable polar contacts for recovery calculation, its subsequent priority should be further evaluated on the basis of interaction stability during molecular dynamics simulation.

The Tier B group consisted of buried 0 model 7 sample 1, buried 7 model 9 sample 4, buried 7 model 2 sample 4, and buried 4 model 0 sample 9. These structures retained partial support for the intended quercetin-recognition mode but were weaker than the Tier A candidates in terms of pose recovery, total contact preservation, or polar-interaction support. Of particular note, buried 0 model 7 sample 1 achieved an affinity of −8.745 kcal mol⁻¹, an in-place RMSD of 1.0008 Å, and complete recovery of reference polar contacts, supporting its use as a priority backup candidate for subsequent dynamic validation.

Focused docking and design-pose recovery analysis identified five Tier A quercetin- binding structures with favorable integrated properties. Among them, buried 1 model 0 sample 8 and buried 5 model 6 sample 8 originated from the strictly selected group, further supporting their structural consistency across both back-prediction and ligand- redocking stages. In addition, buried 6 model 3 sample 2 and buried 4 model 3 sample 8 originated from the conditionally selected group but displayed strong docking-pose recovery characteristics, indicating that focused docking can reveal promising candidates not fully distinguished during the preceding structural screening stage.

Overall, focused molecular docking and reference-pose recovery analysis progressively reduced the candidate sets derived from integrated structural screening. For capsaicin, 19 candidates completed formal docking. For (4R)-limonene, 48 candidates completed docking, yielding seven Tier A structures. For quercetin, 30 candidates completed docking, yielding five Tier A structures. Across all three ligands, candidates with the most favorable Vina scores did not necessarily show the strongest recovery of the design-state ligand pose or contact network. Therefore, reference-pose recovery, ligand displacement, contact preservation, and steric compatibility provided a more informative basis than docking affinity alone for prioritizing candidate binder structures for molecular dynamics simulation.

### 3.7 Molecular dynamics validation of the representative 4DY-binding candidate

To further evaluate the dynamic stability of the representative 4DY-binding protein candidate and its ability to maintain ligand association, partial 5 model 4 sample 5 was subjected to 100-ns molecular dynamics simulations (Figure 5). The representative complex structure showed that 4DY was accommodated within a defined pocket formed by the helical binder scaffold, suggesting that the designed candidate provided a spatially organized environment for ligand recognition.

**Figure 5.**
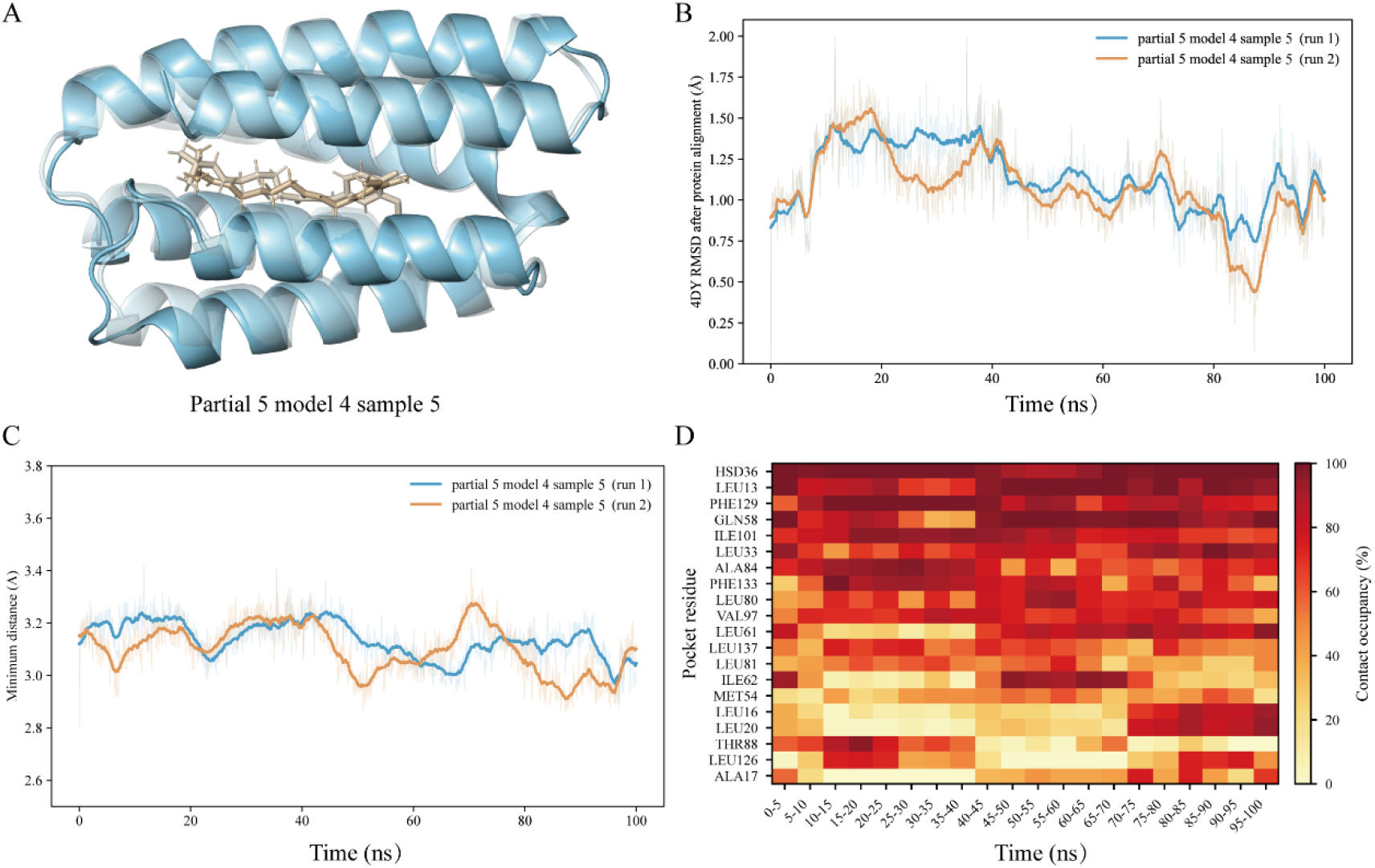
Molecular dynamics-based stability assessment of the 4DY–designed binder complex. Note: A, Structural comparison of the 4DY–binder complex at the initial state (0 ns) and final state (100 ns) of the simulation. B, Heavy-atom RMSD of 4DY calculated after protein-backbone alignment, which was used to evaluate the positional stability of 4DY relative to the binding pocket. Pale curves represent the raw RMSD data, whereas solid curves represent the moving-average-smoothed profiles. C, Minimum distance between 4DY heavy atoms and protein heavy atoms during the 100-ns simulation. The distance remained within a close-contact range overall, suggesting that 4DY maintained persistent contact with the designed binding pocket during the simulation. D, Contact-occupancy heatmap of pocket residues interacting with 4DY across the 100-ns simulations. Residues with high contact occupancy indicate persistent contributions to ligand recognition and pocket stabilization.

Across the two simulation runs, the 4DY heavy-atom RMSD calculated after protein-backbone alignment remained within a limited range. Although local fluctuations were observed during the trajectories, the RMSD curves did not show a continuous increase, indicating that 4DY underwent local pose adjustment rather than persistent displacement from the binding pocket. The two runs displayed broadly similar RMSD trends, while showing local differences at several time intervals, suggesting that the binding mode was generally reproducible but allowed moderate ligand rearrangement within the pocket.

The minimum distance between 4DY and the protein remained within a narrow close-contact range throughout the simulations. No long-term increase in the minimum distance was observed, indicating that 4DY maintained persistent contact with the binder during the 100-ns trajectories. This result was consistent with the ligand RMSD analysis and further supported the absence of obvious ligand escape from the designed binding site.

The pocket-residue contact-occupancy heatmap further identified residues that contributed persistently to ligand recognition. Several residues, including HSD36, LEU13, PHE129, GLN58, ILE101, LEU33, ALA84, PHE133, LEU80, VAL97, and LEU61, showed high contact occupancy across multiple time windows. These residues may form a major interaction interface for 4DY and contribute to pocket stabilization through hydrophobic packing, spatial confinement, and local polar interactions. Collectively, the RMSD, minimum-distance, and contact-occupancy analyses support partial 5 model 4 sample 5 as a dynamically stable 4DY-binding candidate.

## 4 Discussion

In this study, we established a staged computational protein design and screening workflow for plant-derived small molecules and evaluated its applicability to the design of natural-product-recognizing proteins using capsaicin, (4R)-limonene, and quercetin as model ligands with distinct structural features and functional attributes. These molecules are associated with pungent flavor, volatile aroma, and flavonoid-related bioactive constituents in characteristic agricultural products and medicinal–edible resources from the Guanzhong and Qinba regions of Shaanxi Province. Rather than simply expanding the number of generated candidates, this study focused on progressively identifying structures that retained the intended ligand-recognition logic across different computational levels. JSON-based quality assessment of RFdiffusion3 outputs was effective in excluding initial backbones with evident chain breaks, steric conflicts, or unfavorable global geometry; however, favorable overall structural quality did not necessarily indicate that a candidate had formed a suitable binding pocket around the target ligand. Indeed, for capsaicin, (4R)-limonene, and quercetin, some structures with favorable initial global rankings were no longer among the optimal pocket-forming candidates after ligand-centred local evaluation. Therefore, PyMOL-based inspection of the local ligand environment was not merely supplementary to initial structural screening, but was required to determine whether candidate backbones truly satisfied the intended buried or partial-exposure design constraints. More broadly, RFdiffusion3[20], LigandMPNN[14], and RoseTTAFold3[15] focus on complementary but distinct aspects of the design problem, namely backbone generation, sequence compatibility with the local structural environment, and predicted folded conformation, respectively. No single output from these stages can fully determine whether a candidate possesses a reasonable initial pocket, an implementable amino acid sequence, a back-predicted structure that preserves the intended conformation, and an ability to recover the designed small-molecule recognition mode. We therefore implemented a progressively convergent evaluation strategy that shifted candidate prioritization from single-score ranking toward integrated assessment of design consistency.

This multi-stage screening strategy led to a stepwise convergence of candidate structures for all three ligands. At the RFdiffusion3 backbone-generation stage, 100 initial candidate backbones were generated for capsaicin, 100 for (4R)-limonene, and 200 for quercetin. Following initial structural quality screening based on JSON outputs, 86, 95, and 154 backbones, respectively, were advanced to PyMOL-based local pocket evaluation. Ligand-centred PyMOL screening subsequently retained 31 capsaicin partial-pocket backbones, 75 (4R)-limonene buried-pocket backbones, and 56 pocket-qualified quercetin backbones. For each of these retained backbones, LigandMPNN was used to design 10 candidate sequences, followed by RoseTTAFold3-based structural back-prediction, resulting in 310, 750, and 560 protein structures for capsaicin, (4R)-limonene, and quercetin, respectively. By integrating predicted stability, solubility and usability, pocket RMSD, recovery of original ligand contacts, interface sequence recovery, and spatial compatibility with the projected protein–ligand clashes, we ultimately identified 31 capsaicin-, 54 (4R)-limonene-, and 32 quercetin-binding candidate structures for subsequent molecular docking analysis. Among the 31 representative capsaicin structures, 25 showed no detectable ligand steric clashes, and 16 simultaneously exhibited clash-free geometry and high projected contact recovery. For (4R)-limonene, 28 structures were classified as strictly selected and 26 as conditionally selected. For quercetin, 11 structures were strictly selected and 21 were conditionally selected. Together, these findings indicate that acceptable initial backbone quality only provides the structural basis for further evaluation; preservation of pocket geometry after sequence design, compatibility with the intended ligand pose, and recovery of the designed contact pattern are required to determine whether a structure is suitable for downstream validation.

The differences observed among the three ligand systems further suggest that pocket- design strategies for natural-product recognition should be adapted to the physicochemical characteristics of individual ligands. Capsaicin contains an aromatic recognition region, a flexible hydrophobic chain, and functional groups capable of participating in polar interactions[24]. A partial-exposure strategy was therefore appropriate for enclosing its principal hydrophobic recognition region while retaining reasonable accessibility of selected polar groups. In contrast, (4R)-limonene is a compact and highly hydrophobic monoterpene lacking conventional hydrogen-bond donors or acceptors[25], making a buried-ligand strategy centred on hydrophobic enclosure consistent with its likely recognition requirements. The quercetin[26] results were particularly informative: under the design conditions used in this study, 55 of 75 buried-mode candidates passed PyMOL-based local pocket evaluation, whereas only 1 of 79 partial-mode candidates satisfied the screening criteria. This marked difference suggests that, for quercetin, which possesses a rigid planar aromatic scaffold and multiple polar groups, structures that more extensively enclose the ligand scaffold while organizing an appropriate internal polar interaction environment may be more readily generated than structures that reserve a relatively large solvent-exposed region. Because the two quercetin design tasks were not identical with respect to atom- conditioning schemes and designed length ranges, these findings should not be interpreted as demonstrating the universal superiority of the buried-ligand strategy under all conditions. Rather, they indicate that, under the parameter settings and screening criteria applied here, the buried-ligand configuration was more effective in generating quercetin pocket candidates suitable for subsequent evaluation. This observation highlights a broader design principle: protein recognition of natural-product small molecules is unlikely to be addressed by a universal pocket template, but instead requires ligand-specific constraints informed by hydrophobicity, rigidity, polarity distribution, and the spatial organization of potential interactions.

Following structural back-prediction and multi-parameter screening, focused molecular docking and reference-pose recovery analysis were further used to evaluate whether each target ligand could re-enter the candidate pocket while retaining the binding mode predefined during the design stage. In total, 19 capsaicin, 48 (4R)-limonene, and 30 quercetin candidate structures completed formal molecular docking analysis. A common and methodologically important observation across all three ligand systems was that candidates with more favorable predicted binding energies did not necessarily show better recovery of the reference binding pose. In other words, a more favorable docking affinity could correspond to an alternative low-energy binding orientation rather than the recognition mode intended by the original design. This distinction is especially important for *de novo*-designed small-molecule pockets, because the objective is not merely to allow a ligand to occupy a protein cavity, but to preserve a defined spatial recognition logic. Accordingly, integrated evaluation of docking energy, pose displacement, recovery of original contacts, formation of new contacts, and spatial compatibility provides a more appropriate basis for prioritizing candidate structures than affinity-based ranking alone. This finding suggests a potentially generalizable principle for post-design screening of artificial small-molecule-binding proteins: when a design workflow begins from a predefined pocket and ligand pose, consistency with the designed pose should be treated as an evaluation dimension at least as important as, and in some circumstances more informative than, predicted docking energy.

From an application perspective, the significance of this study is not that it replaces established chemical analytical technologies such as high-performance liquid chromatography or liquid chromatography–mass spectrometry, but rather that it provides potential engineerable biorecognition modules for natural-product small- molecule detection and enrichment. Chemical analytical approaches remain essential for the accurate identification and quantification of target constituents in complex samples, but they do not directly produce molecular recognition elements that can be further incorporated into capture systems, competitive detection formats, or sensor interfaces[27, 28]. Artificially designed protein binders, by contrast, potentially offer encodable sequences, optimizable structures, engineerable recognition surfaces, and compatibility with downstream signal-output modules[13]. Therefore, if the candidate structures generated for capsaicin, (4R)-limonene, and quercetin are subsequently validated to possess sufficient affinity and selectivity, they may provide a basis for developing biorecognition tools for assisting pungency-related evaluation, detecting citrus aroma-associated constituents, and enriching or analysing flavonoid-related bioactive components. For example, candidate capsaicin-binding proteins could provide molecular starting points for rapid detection of pungent constituents in chili products such as Shaanxi-style *youpo lazi*; candidate (4R)-limonene binders could support studies of volatile constituent recognition in citrus and other aromatic resources; and candidate quercetin binders could be further investigated for flavonoid detection and enrichment in *Cornus officinalis*, *Epimedium* species, and other plant-derived medicinal–edible resources. Importantly, all of these applications depend on subsequent expression, binding, and selectivity validation and should therefore be regarded as potential directions opened by the present study rather than as demonstrated functional outcomes.

This study has several limitations. *De novo* design of a small-molecule-binding protein compresses a discovery process that once took years into just a few hours. This is an idealistic form of generation—it has the potential to create synthetic peptides that surpass natural antibodies, but it also risks producing sequences that are not sufficiently realistic. Therefore, a realistic approach to mining is also required, to identify from an extremely vast potential sequence space the small subset of peptides with the highest activity and lowest toxicity. From computational prediction to practical application, there remain real-world gaps involving data, toxicity, modified peptides, and *in vivo* validation. This study has not yet been validated through recombinant expression, *in vitro* affinity measurements, discrimination of structurally related ligands, or sensor performance testing. Consequently, the structures identified here should be regarded as candidates prioritized for experimental validation rather than as functionally confirmed protein binders. Accordingly, the structures identified here should be regarded as candidates with priority for experimental validation rather than as functionally confirmed protein binders. Second, the docking evaluation used the RFdiffusion3 design-state ligand position as a reference. This approach is advantageous for determining whether a candidate preserves the intended recognition mode, but it cannot independently demonstrate the true binding preference of a protein in an unconstrained setting, nor can it directly establish selectivity against structurally related natural-product molecules. Future studies should therefore incorporate cross-ligand docking, structurally related analogue or decoy-ligand evaluation, and competitive binding experiments to assess candidate selectivity. Third, owing to current computational resource limitations, only one representative high-priority docked capsaicin–binder complex was selected for exploratory molecular dynamics simulations. Consequently, the dynamic analyses primarily provide an initial assessment of whether complexes obtained through this workflow can retain basic binding-pocket and ligand-position stability under dynamic conditions, rather than a sufficient basis for final ranking of all candidates. Future work should prioritize systematic, replicated molecular dynamics simulations of candidates showing the strongest pose-recovery and contact- preservation characteristics, followed by experimental assessment of protein expression, thermal stability, and binding properties using techniques such as microscale thermophoresis, surface plasmon resonance, biolayer interferometry, or isothermal titration calorimetry.

In conclusion, this study not only identified candidate binding structures for capsaicin, (4R)-limonene, and quercetin, but also established a staged computational framework spanning ligand-constraint-driven backbone generation, local pocket screening, sequence design, structural back-prediction, reference-pose recovery evaluation, and exploratory analysis of dynamic stability. Based on the retained candidate structures, an initial candidate protein-binder resource library for these three representative natural-product small molecules was organized to include candidate identifiers, target ligands, amino acid sequences, protein lengths, design modes, predicted structures, docked poses, key contacting residues, docking scores, structural validation metrics, and priority classifications, with recombinant expression, binding-affinity, and selectivity data to be progressively incorporated in future studies. It provides testable candidate resources for the future development of biorecognition and biosensing tools that may assist food-flavor evaluation, natural-product constituent detection, and bioactive-component enrichment. In parallel, the hierarchical screening logic and automated analysis scripts developed in this study may be extended to additional plant-derived small molecules and iteratively refined through experimental feedback, thereby supporting the gradual establishment of a design–screen–validate–redesign cycle for the development of natural-product-recognizing proteins and providing a reusable computational starting point for programmable biorecognition of complex natural-product resources.

## Data and model availability statement

The dataset and candidate binder models generated in this study are available in the GitHud repository: https://github.com/ZhuYaojun1/Design-and-prioritization-of-candidate-protein-binders-for-capsaicin-4R--limonene-and-quercetin

## Authors’ contributions

YZ: Conceptualization, Methodology, Investigation, Formal analysis, Visualization, Writing – original draft. XZ: Conceptualization, Supervision, Funding acquisition, Project administration, Writing – review & editing. All authors read and approved the final manuscript.

## References

1. Mullowney MW, Duncan KR, Elsayed SS, Garg N, van der Hooft JJJ, Martin NI, Meijer D, Terlouw BR, Biermann F, Blin K et al: Artificial intelligence for natural product drug discovery. Nat Rev Drug Discov 2023, 22(11):895–916. doi: 10.1038/s41573-023-00774-7

2. Vitale GA, Geibel C, Minda V, Wang M, Aron AT, Petras D: Connecting metabolome and phenotype: recent advances in functional metabolomics tools for the identification of bioactive natural products. Nat Prod Rep 2024, 41(6):885–904. doi: 10.1039/d3np00050h

3. Xi C, Diao J, Moon TS: Advances in ligand-specific biosensing for structurally similar molecules. Cell Syst 2023, 14(12):1024–1043. doi: 10.1016/j.cels.2023.10.009

4. Zhao Y, Hao H, Geng X, Jia M, Zhang X, Wang M, Yang Y, Li Y, Wang S, Zheng X: The historical evolution and research progresses of three Qinba characteristic Chinese medicinal materials. Journal of Northwest University (Natural Science Edition) 2024, 54(5):767–784. doi: 10.16152/j.cnki.xdxbzr.2024-05-002

5. Chaisupa P, Wright RC: State-of-the-art in engineering small molecule biosensors and their applications in metabolic engineering. SLAS Technol 2024, 29(2):100113. doi: 10.1016/j.slast.2023.10.005

6. Snoek T, Chaberski EK, Ambri F, Kol S, Bjørn SP, Pang B, Barajas JF, Welner DH, Jensen MK, Keasling JD: Evolution-guided engineering of small-molecule biosensors. Nucleic Acids Res 2020, 48(1):e3. doi: 10.1093/nar/gkz954

7. Lee GR, Pellock SJ, Norn C, Tischer D, Dauparas J, Anishchenko I, Mercer JAM, Kang A, Bera AK, Nguyen H et al: Small-molecule binding and sensing with a designed protein family. Nat Commun 2026, 17(1). doi: 10.1038/s41467-026-70953-8

8. Juárez-Contreras R, Mota-Carrillo E, Piedra-Ramírez A, Farías-Sánchez D, González-Ramírez R, Morales-Lázaro SL: Capsaicin: beyond TRPV1. Front Nutr 2025, 12:1594742. doi: 10.3389/fnut.2025.1594742

9. He J, Qin Z, Liu K, Li X, Kou Y, Jin Z, He R, Hong M, Xiong B, Liao L et al: Volatile metabolomics and transcriptomics analyses provide insights into the mechanism of volatile changes during fruit development of ‘Ehime 38’ (Citrus reticulata) and its bud mutant. Front Plant Sci 2024, 15:1430204. doi: 10.3389/fpls.2024.1430204

10. Cho J, Kim K, Lee J-G: Quercetin in food: structure, biosynthesis, toxicity, analytical method, occurrence and risk assessments. Applied Biological Chemistry 2025, 68(1):90. doi: 10.1186/s13765-025-01063-0

11. Broeckling CD, Beger RD, Cheng LL, Cumeras R, Cuthbertson DJ, Dasari S, Davis WC, Dunn WB, Evans AM, Fernández-Ochoa A et al: Current Practices in LC-MS Untargeted Metabolomics: A Scoping Review on the Use of Pooled Quality Control Samples. Anal Chem 2023, 95(51):18645–18654. doi: 10.1021/acs.analchem.3c02924

12. Sadybekov AV, Katritch V: Computational approaches streamlining drug discovery. Nature 2023, 616(7958):673–685. doi: 10.1038/s41586-023-05905-z

13. Yang W, Wang S, Lee GR, Zhang JZ, Courbet A, Juergens D, Wang X, Schlichthaerle T, Abedi M, Ragotte R et al: The past, present and future of de novo protein design. Nature 2026, 652(8112):1139–1152. doi: 10.1038/s41586-026-10328-7

14. Dauparas J, Lee GR, Pecoraro R, An L, Anishchenko I, Glasscock C, Baker D: Atomic context-conditioned protein sequence design using LigandMPNN. Nat Methods 2025, 22(4):717–723. doi: 10.1038/s41592-025-02626-1

15. Corley N, Mathis S, Krishna R, Bauer MS, Thompson TR, Ahern W, Kazman MW, Brent RI, Didi K, Kubaney A et al: Accelerating Biomolecular Modeling with AtomWorks and RF3. bioRxiv 2025. doi: 10.1101/2025.08.14.670328

16. Errington D, Schneider C, Bouysset C, Dreyer FA: Assessing interaction recovery of predicted protein-ligand poses. J Cheminform 2025, 17(1):76. doi: 10.1186/s13321-025-01011-6

17. Eberhardt J, Santos-Martins D, Tillack AF, Forli S: AutoDock Vina 1.2.0: New Docking Methods, Expanded Force Field, and Python Bindings. J Chem Inf Model 2021, 61(8):3891–3898. doi: 10.1021/acs.jcim.1c00203

18. Lemkul JA: Introductory Tutorials for Simulating Protein Dynamics with GROMACS. J Phys Chem B 2024, 128(39):9418–9435. doi: 10.1021/acs.jpcb.4c04901

19. Westbrook JD, Shao C, Feng Z, Zhuravleva M, Velankar S, Young J: The chemical component dictionary: complete descriptions of constituent molecules in experimentally determined 3D macromolecules in the Protein Data Bank. Bioinformatics 2015, 31(8):1274–1278. doi: 10.1093/bioinformatics/btu789

20. Butcher J, Krishna R, Mitra R, Brent RI, Li Y, Corley N, Kim PT, Funk J, Mathis S, Salike S et al: De novo Design of All-atom Biomolecular Interactions with RFdiffusion3. bioRxiv 2025. doi: 10.1101/2025.09.18.676967

21. Thumuluri V, Martiny HM, Almagro Armenteros JJ, Salomon J, Nielsen H, Johansen AR: NetSolP: predicting protein solubility in Escherichia coli using language models. Bioinformatics 2022, 38(4):941–946. doi: 10.1093/bioinformatics/btab801

22. Gasteiger E, Hoogland C, Gattiker A, Duvaud Se, Wilkins MR, Appel RD, Bairoch A: Protein identification and analysis tools on the ExPASy server. In: The proteomics protocols handbook. Springer; 2005: 571–607.

23. Van Der Spoel D, Lindahl E, Hess B, Groenhof G, Mark AE, Berendsen HJ: GROMACS: fast, flexible, and free. J Comput Chem 2005, 26(16):1701–1718. doi: 10.1002/jcc.20291

24. 4DY [https://www.rcsb.org/ligand/4DY]

25. 9IR [https://www.rcsb.org/ligand/9IR]

26. QUE [https://www.rcsb.org/ligand/QUE]

27. Chen L, Xu S, Li J: Recent advances in molecular imprinting technology: current status, challenges and highlighted applications. Chem Soc Rev 2011, 40(5):2922–2942. doi: 10.1039/c0cs00084a

28. Turner AP: Biosensors: sense and sensibility. Chem Soc Rev 2013, 42(8):3184–3196. doi: 10.1039/c3cs35528d

